# Performance Test of the QNome Nanopore Sequencer for Arthropod DNA Barcoding

**DOI:** 10.64898/2026.04.29.721586

**Authors:** Ken A Thompson, Sean WJ Prosser, Robin M Floyd, Saeideh Jafarpour, Emine Ozsahin, Paul DN Hebert

## Abstract

Nanopore sequencers have the potential to liberate DNA sequencing from centralized core facilities to distributed analytical nodes. Until recently, Oxford Nanopore Technologies (ONT) had been the sole manufacturer of a portable nanopore sequencer, but analogous platforms are in production. Nanopore sequencers from Qitan Technology (QT) are widely used in China but have been unavailable outside that nation and have not been subject to rigorous, independent performance testing. Enabled by early access to QT’s least expensive sequencer and flow cell, the QNome-3841 and QCell-384, we tested whether they could generate accurate DNA barcodes cost-effectively. In several tests involving amplicon pools from 96 to 9,120 specimens, QT recovered valid DNA barcodes from nearly as many specimens (98.5%) as ONT. QT sequences had slightly lower accuracy than their ONT counterparts (median per-read identity 98.04% vs. 99.39%) and QT frequently failed to resolve the correct length of G/C homopolymers. However, barcode sequences from the two platforms were nearly indistinguishable after bioinformatic treatment with the Barcode Inference Pipeline (BIP). QT’s wash kit performed well, enabling a QCell to sequence eight amplicon pools with negligible carryover (four of >800,000 reads) among eight runs, and a gradual degradation of the flow cell. Its ultra-fast protocol allowed library preparation in a single step that could be completed in under 10 minutes, but it led to a significant reduction in read retention. Once widely available, QT devices will be well-suited to support DNA barcode analysis.

## INTRODUCTION

By coupling its status as the first market entrant with a strong record of innovation, Oxford Nanopore Technologies (ONT) has held a near-monopoly on nanopore sequencing for a decade. Its first sequencer, the MinION, was introduced in 2015 with the goal of making it possible for anyone to sequence anything, anywhere. Its portability, low cost, long reads, and USB power requirement attracted immediate attention. While initial uptake was slowed by low sequence quality, a key technological innovation in 2022 greatly improved read fidelity (Zhang et al., 2023). Over the decade, ONT has diversified its sequencing platforms and flow cells, increasingly with a view towards supporting the sequencing of whole genomes. This focus has led it to emphasize high-throughput sequencing platforms, to step back from certain products, and to escalate prices. For example, in 2026 ONT announced both the discontinuation of the Flongle flow cell (Fig. S1) and its least expensive sequencer capable of running the high-throughput PromethION flow cell (P2 Solo; Fig. S2). ONT’s shifting product line has created opportunities for new entrants into the nanopore marketplace.

Several firms now produce nanopore sequencers that address differing applications and leverage varying underlying technological approaches. Roche’s SBX nanopore sequencer is a high-cost instrument, suited for deployment in core facilities that are sequencing whole genomes at large scale (Kokoris et al., 2025). At least five Chinese companies—AxBio, BGI, Genvida, RH Genetech and Qitan Technology—are developing nanopore sequencers for a spectrum of applications (Whiteford, 2022a). Among the firms developing portable sequences, Qitan (hereafter ‘QT’), is considered by industry experts to be the most prominent firm among them (Whiteford, 2022b). QT manufactures nanopore sequencers, flow cells, associated consumables and software (https://www.qitantech.com/). Its QNome-3841 (hereafter ‘QNome’; Figure 1) sequencer is a close equivalent to the MinION both in cost (Qitan Tech, pers. comm.) and stated capabilities (https://www.qitantech.com/products/sequencer/platform/QNome), but it is not currently widely available. QT offers higher-throughput instruments and flow cells via the QPursue and QPinnacle series (Dang et al., 2026), but they are not evaluated here or discussed further.

**Figure 1.**
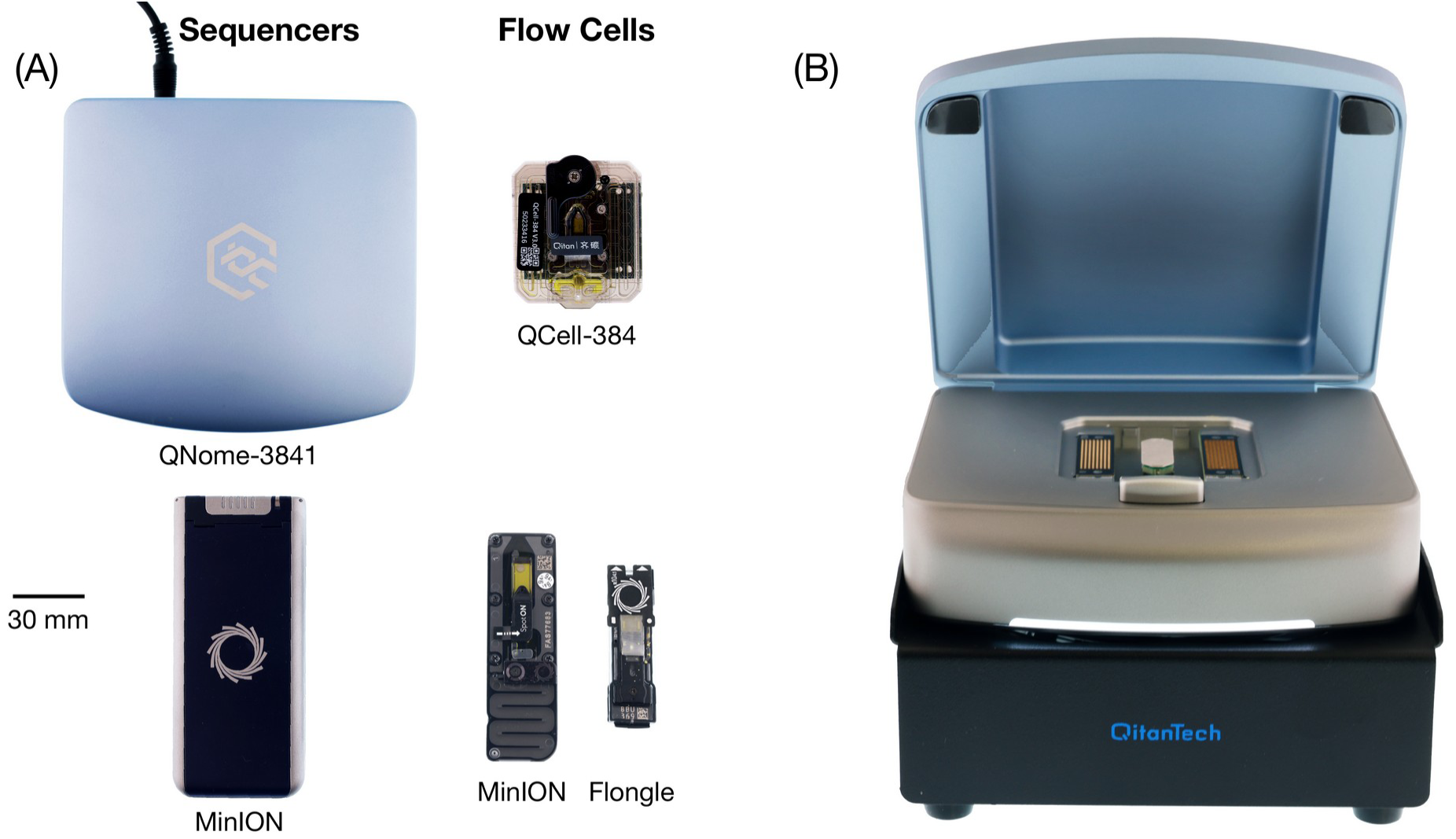
Photographs of the QT and ONT sequencers and flow cells used in this study. (A) Top- down view of sequencers (left) and flow cells (centre); scale bar applies to all. (B) Photograph of QNome sequencer on the external fan with the lid open. The insertion point for a QCell is visible.

To generate sequence data, all nanopore sequencers require consumable flow cells which can be run once or washed for reuse. In the case of the MinION, there are two options: the soon-to-be-discontinued Flongle ($210 USD per run, including reagents) which generates 2.8 Gb and the MinION ($960 USD) which generates 48 Gb. By comparison, the QNome only employs the QCell-384 ($280 USD; hereafter ‘QCell’; Figure 1) which generates 9 Gb. Considering these three options, sequencing costs per Gb vary from $75 for Flongle to $20 for MinION and $30 for QCell. The QCell and MinION flow cells have a major advantage over the Flongle as they can be reused several times while the Flongle is single use.

Targeted amplicon sequencing underpins the DNA-based identification systems which have gained broad adoption by the biodiversity science community (CBOL Plant Working Group, 2009; Hebert et al., 2003; Schoch et al., 2012). DNA barcoding, and related approaches such as organismal and eDNA metabarcoding, are widely used in biodiversity assessment (Yu et al., 2012). Matching amplicon sequences to a reference library enables rapid and inexpensive inventories (Morinière et al., 2019). Portable sequencers capable of on-site sequencing are increasingly valued under the Nagoya Protocol and in communities where data sovereignty is prioritized (Gygax et al., 2025). ONT’s MinION sequencer and flow cell were quickly adopted for DNA barcoding outside of core lab facilities (Pomerantz et al., 2018) and are well-validated by the community (Hebert et al., 2025). The extensive use of MinION for barcoding motivated the development of software designed to support this use case (Srivathsan et al., 2018, 2021, 2024). Because of their low cost, Flongles were transformative for barcoding, so they were quickly adopted (Hebert et al., 2025; Koblmüller et al., 2024; Srivathsan et al., 2024). However, ONT’s shift away from low-priced sequencers and flow cells creates an incentive to investigate other platforms. Based on analytical specifications and cost, the QNome sequencer and QCell appear well positioned to support amplicon sequencing, but there has been no independent test of their performance. In fact, only one study has assessed the capacity of a QT sequencer to support specimen identification through amplicon sequencing (Zhao et al., 2026). That study, which involved QT-affiliated authors, sequenced 60 samples for 12S rRNA and reported 99.34% identity to “gold-standard” Sanger sequences with most of the sequence divergence reflecting errors in homopolymer regions. Compared to that study, ours is larger by two orders of magnitude and we also are independent of QT. We also evaluate QT’s ultra-fast library preparation kit and the wash kit. In the present study, we examine the capacity of QNome to recover DNA barcodes from the most diverse lineage of terrestrial life: arthropods. Specifically, we tested its performance by sequencing the same amplicon pools on both the QNome and MinION. We sequenced a pool of nearly 10K specimens to compare QCell to MinION (Experiment 1) and a second pool of 2K specimens to compare it with Flongle (Experiment 2). We also evaluated the performance of the QCell over multiple washes (Experiment 3) and tested its ultra-fast (<10-minute) library preparation kit (Experiment 4).

## MATERIALS AND METHODS

Although its products are not generally available outside China, we acquired key items from QT. Specifically, we purchased a QNome-3841 sequencer, 18 QCell-384 flow cells, two library preparation kits, an ultra-fast library preparation kit, three flow cell wash kits, and three DNA sequencing kits. We obtained access to QT products by first sending an inquiry to the main customer intake address on its website and then corresponding with the QT employee in charge of overseas sales. No formal agreements were signed, and all equipment and reagents were used without restriction. No equipment was loaned, and we paid the prices quoted by QT without negotiation.

We used the standard library preparation kit for Experiments 1–3 and the ultra-fast kit for Experiment 4; all runs used QT’s DNA Sequencing Kit to load the flow cell. PCR pools were prepared for sequencing on QT and ONT following manufacturer’s protocols and reaction volumes. All ONT sequencing used a MinION Mk-1d sequencer. The QNome is roughly 2.5× larger than the MinION (Figure 1) and requires access to 110V–220V power and an external fan, while the MinION is stand- alone and USB-powered.

All experiments employed similar protocols bar the unique aspects detailed below. We also describe the bioinformatics and other downstream analyses used to process and evaluate data quality.

Except where noted, DNA extraction and PCR followed standard protocols at QT sequencing used QT software (QPreasy v. 3.4.0) to interact with the QNome. It was installed on an Alienware Aurora R15 AMD desktop computer running Ubuntu 22.04, equipped with 64 GB memory, an AMD® Ryzen 9 7950× 16-core processor × 32, and an NVIDIA GeForce RTX 4090 GPU. Basecalling was performed live using QPreasy’s built in basecalling model: QDSv3.0_QCell384v3.0_QDLEv3.0_NTGv5.1.1T. Full log files—containing complete and granular detail about all software versions and parameters—are available for all runs.

For ONT sequencing, we used MinKNOW v. 25.09.16 to interact with the MinION sequencer. MinKNOW was installed on an Alienware Aurora R13 computer with 96 GB memory, running an Intel Core i9-12900KF × 24 processor and equipped with an NVIDIA GeForce RTX 3080 GPU. Basecalling was performed live using Dorado 7.11.2 with the super-accurate model dna_r10.4.1_e8.2_400bps_sup@v5.2.0, and MinKNOW’s minimum Q-score was set to 10. ONT libraries were prepared with the Ligation Sequencing Kit V14 (SQK- LSK114) and sequenced on R10.4.1 flow cells: a MinION flow cell (FLO-MIN114) in Experiment 1 and a Flongle flow cell (FLO-FLG114) in Experiment 2. Like QT, full log files with comprehensive granular details are available for all ONT runs.

Because different computers were used to control the two sequencers, all durations reported in this text should be viewed as a statement of fact and not for comparison between the devices. Anecdotally, basecalling was not a rate-limiting step for either sequencer. We also note that the three flow cell types used herein— Flongle, MinION, and QCell-384—have different numbers of pores, and differing rates of pore deactivation during runs. Standardizing initial pore count is not possible, so we standardize the comparisons by read count and by enforcing identical minimum quality scores.

### Experiment 1: Performance of QCell and MinION flow cells

This study compared data derived from a QCell with that obtained from a MinION flow cell. Analysis focused on the characterization of an amplicon pool derived from 9,120 specimens mostly collected by Malaise traps (99%; 95 specimens hand- collected) deployed in New South Wales and Tasmania, Australia in 2010, 2011, and 2024. Specimen records are in a dataset (DS-QT96) on BOLD (Ratnasingham et al., 2024) (www.boldsystems.org). Insects predominated (89.4%), followed by arachnids (7.7%; mainly mites) and springtails (2.8%). The main insect orders were Diptera (34.7% of specimens), Lepidoptera (19.7%), Hymenoptera (18.9%), Hemiptera (9.0%) and Coleoptera (4.7%).

After PCR amplification, indexing, and cleanup, aliquots from the PCR pool were used as a basis for library preparation following both ONT and QT protocols. ONT protocols followed those detailed by Floyd et al. (2023) but used the manufacturer’s recommended volumes. QT has not published its protocols, so we give them in full in Supplementary Text S1. The ONT library was sequenced on a MinION flow cell (FLO-MIN114, R10.4.1) with 1,255 initial pores on the pore scan. The QCell had 192 initial pores. Both sequencing runs proceeded for 72 hr.

### Experiment 2: Performance of QCell and Flongle flow cells

This study compared the data derived from a QCell flow cell with that from a Flongle. We selected 24 plates of extracted DNA expected to yield high barcode recovery for PCR amplification for COI and prepared sequencing libraries from them. Specimens were derived from Malaise traps deployed in December 2024 in Muogamarra, NSW, Australia. All specimen records are in a dataset (DS-QT24) on BOLD. Fourteen of these plates (1,330 specimens; 58.3%) were also part of DS- QT96. Insects again predominated (95.4%), led by Lepidoptera (33.8%), Diptera (26.8%), Hymenoptera (12.4%), Hemiptera (12.3%) and Coleoptera (8.6%).

The Flongle library was sequenced first. The Flongle had 55 initial pores and generated 1,047,440 reads over 24 hr before filtering. The QCell library was then analyzed and QPreasy was programmed to stop the run after it reached 1M reads (261 active pores), which required 2 h 50 min. We note that the design—specifically the channel and pore structure—of the Flongle flow cell differs from that of the MinION flow cell (Pugh, 2023), so this experiment is analyzed separately from Experiment 1.

### Experiment 3: Effectiveness of QCell flow cell wash

QCells can be washed to allow their reuse. We tested QT’s washing protocol to determine whether it prevented the carryover of DNA from one run to the next. This study involved sequencing a library derived from one 96-well plate, washing the flow cell, and loading it with a different library for sequencing. This cycle of washing, reloading, and sequencing was repeated for eight QNome runs. All wells in each of the eight plates contained a single species’ template DNA in each well, and the species differed among plates so any carryover would be recognized. Details about the specimens sequenced and DNA extraction method are provided in Supplementary Text S3.

Each PCR plate had the same schema of forward UMIs but a distinct reverse UMI. PCR, library preparation, and sequencing on the QCell employed the same procedure as Experiments 1 and 2. The first three sequencing runs and washes were completed in one day. The fourth through eighth runs were completed in a single day, 23 days after the first runs. QPreasy was programmed to stop after 100K reads for each run. We analyzed each library using the same input parameter files, so all subsequent and prior libraries could be detected in each run. Detection of sequences from prior libraries would indicate that washing the flow cell failed to clear all DNA.

### Experiment 4: Ultra-fast library preparation

A final study evaluated the effectiveness of QT’s ultra-fast library preparation kit. Using the same 24-plate PCR pool as for Experiment 2, we completed the ultra-fast library preparation protocol and then sequenced the library on a new QCell. Qitan’s ultra-fast protocol differs from the main protocol in that it requires just a single 5- min incubation.

The ultra-fast library was analyzed on a different QCell than the standard library because they use different sequencing adapters that QPreasy was programmed to recognize and trim.

The QCell used for Expt 4 initially had 202 pores, ran overnight, and was stopped the following day; it generated 3.85M reads passing the quality filter in 20 h 29 min and was downsampled for analysis.

### Bioinformatics and Analysis of Sequence Quality

Bioinformatic analysis used BIP v1.1.0 (Prosser et al., 2026) with default parameters for COI-5P (see Data Accessibility for input files). BIP is the bioinformatics pipeline employed by the CBG’s core facility to support the barcode analysis of roughly 3 million individual specimens annually. BIP takes raw sequencer output, a reference library, and a parameters file, and returns tabular data with target and non-target barcode sequences together with their associated taxonomic assignments.

Briefly, BIP orients sequencing reads, assigns them to primer pairs, and discards reads outside a user-specified (here 550–950 bp) length window while UMIs and primers are still attached. It then demultiplexes reads to wells by their forward and reverse UMIs and trims UMIs and primers using cutadapt (Martin, 2011). Within each well, BIP clusters reads into Operational Taxonomic Units (OTUs) —similarity-based sequence groups serving as taxon proxies—with vsearch (Rognes et al., 2016) in two passes, a primary clustering at a specified (here 95%) identity followed by a secondary clustering at a higher specified identity (here 97%), retaining OTUs supported by a minimum number of reads, here five. It builds a consensus sequence for each OTU from a homopolymer-aware multiple sequence alignment of distinct reads, weighted by read count, generated with MAFFT (Katoh & Standley, 2013), and removes chimeras using vsearch’s *de novo* implementation of UCHIME (R. C. Edgar et al., 2011). OTUs are assigned taxonomy with SINTAX (R. Edgar, 2016) against a reference library, here BOLDistilled (Prosser et al., 2025), using a confidence threshold (here 0.6), and are matched to BINs—a COI specific OTU system used for Animalia (Ratnasingham & Hebert, 2013)—by global alignment at a specified identity, here 97%. A decision tree then designates at most one OTU per specimen as the target barcode (Supplementary Text S2). We used the parallel implementation of SINTAX for this paper, which is non- deterministic but faster than the single-core default.

The January 2026 BOLDistilled COI library was used in this study (Prosser et al., 2025). Input files for BIP are provided as supplementary files. Downstream analysis of BIP output used R (R Core Team, 2024) with functions in the tidyverse (Wickham et al., 2019). A full description of BIP’s workflow is given in Supplementary Text S2.

BIP has been refined through long experience with COI-5P and five years of experience with nanopore sequencing. This experience has shown that sequencing errors are most common in homopolymer regions with contiguous runs of four or more identical bases. BIP has an error correction procedure that compares all OTUs against a large dataset of reference COI sequences and either adds 1–2 ambiguous (‘N’) bases in homopolymer regions when their omission would result in a 1–2 bp alignment gap, or deletes 1–2 bases from the homopolymer if they would create a gap in the reference sequence.

Assessments of data quality were based on specimens that returned valid barcode sequences for COI as defined by five criteria:

1. Length from 640–670 bp
2. No stop codons
3. No frameshift indels
4. Fewer than 1% ambiguous bases (i.e., 6 or fewer for a 658 bp amplicon)
5. Match to taxonomy of specimen at the order level.

The same input BIP parameters file was used when comparing ONT and QT runs. For the wash experiment, we used the same parameters file—containing all eight plates—for each of the eight runs. This allowed any specimen from the eight runs to be detected during the analysis of each of the eight runs. The same file was also used for the ultra-fast run as was used for Experiment 2 (Flongle vs. QCell).

We evaluated the relationship between GC content and ambiguous bases, as well as the accuracy of QT reads. Each valid 658 bp barcode was divided into all 653 overlapping 6 bp windows, and each window was assigned to a GC bin by its number of G or C bases (an N counts as neither; 0–1 and 5–6 G/C were pooled). For each bin and platform, the percentage of ambiguous bases was the number of Ns in that bin’s windows divided by all bases in those windows. 95% confidence intervals were obtained from 2,000 bootstrap resamples of barcodes within each platform; barcodes rather than windows were resampled because overlapping windows from one barcode are not independent.

To evaluate read accuracy, we employed several procedures. First, we used the procedure from Hebert et al. (2025) with minor modifications. This analysis used all data for specimens in Experiment 1 where a reference sequence from a PacBio Sequel II instrument was available, and where the ONT and QT datasets had an identical consensus sequence matching the PacBio sequence, with ≥10 reads per platform. Analyses restricted to 5,399 658 bp sequences are presented in the main text for simplicity, and the same analysis for Experiment 2 is given in Fig. S3. Briefly, we first identified the raw reads (UMI- and primer-trimmed) that contributed to each final OTU. Next, we randomly sampled between two and ten reads for each OTU and produced their consensus sequence. This procedure was repeated three times for each read depth, and the median of the three distances was determined. We then computed the sequence distance from each consensus sequence to the PacBio reference sequence. Any distance from the reference was interpreted as an error. We also evaluated estimated per-read accuracy directly by aligning each read to its specimen’s reference sequence with minimap2 (Li, 2018) and evaluating substitutions, insertions, and deletions using the same set of 5,399 specimens analyzed above.

## RESULTS

We begin by summarizing the results from the four experiments before both evaluating the impact of GC content on error rate and documenting read accuracy using data from Experiment 1. We remind the reader that input read counts were standardized between units of comparison as reported in the main text. Full runs are analyzed in the supplementary information (Tables S1–S3) which also provides over 15 summary metrics for each sequencing run, including cost breakdowns (Tables S4–S5; Fig. S4).

### Experiment 1: Performance of QCell and MinION flow cells

The QCell generated 3.1M reads suitable for analysis while the MinION generated 21.1M (Table S4). After downsampling the MinION results to 3.1M reads to enable direct comparison, we found ONT generated valid barcodes for 8,398 (92.1%) specimens while QT generated valid barcodes for 8,249 (90.4%) (Table 1; Figure 2A). A relatively small number of specimens had barcodes recovered only from QT (28 vs. 177 only from ONT); the majority of specimens lacking barcode had no results for either platform (694) (Table S6; Fig. S5A). The majority (74.6%) of ONT and QT sequences were identical, with many additional sequences being identical apart from Ns (21.8%; Table S7; Fig. S5B). BIN and taxonomic assignments were identical for nearly all specimens where both platforms returned a barcode sequence (98.9% and 99.5%, respectively; Table S7; Fig. S5B). Analyzing all 21.1M MinION reads raised ONT recovery only slightly, to 8,528 (93.5%), so a sevenfold increase in reads added 130 barcodes (Table S3).

**Figure 2.**
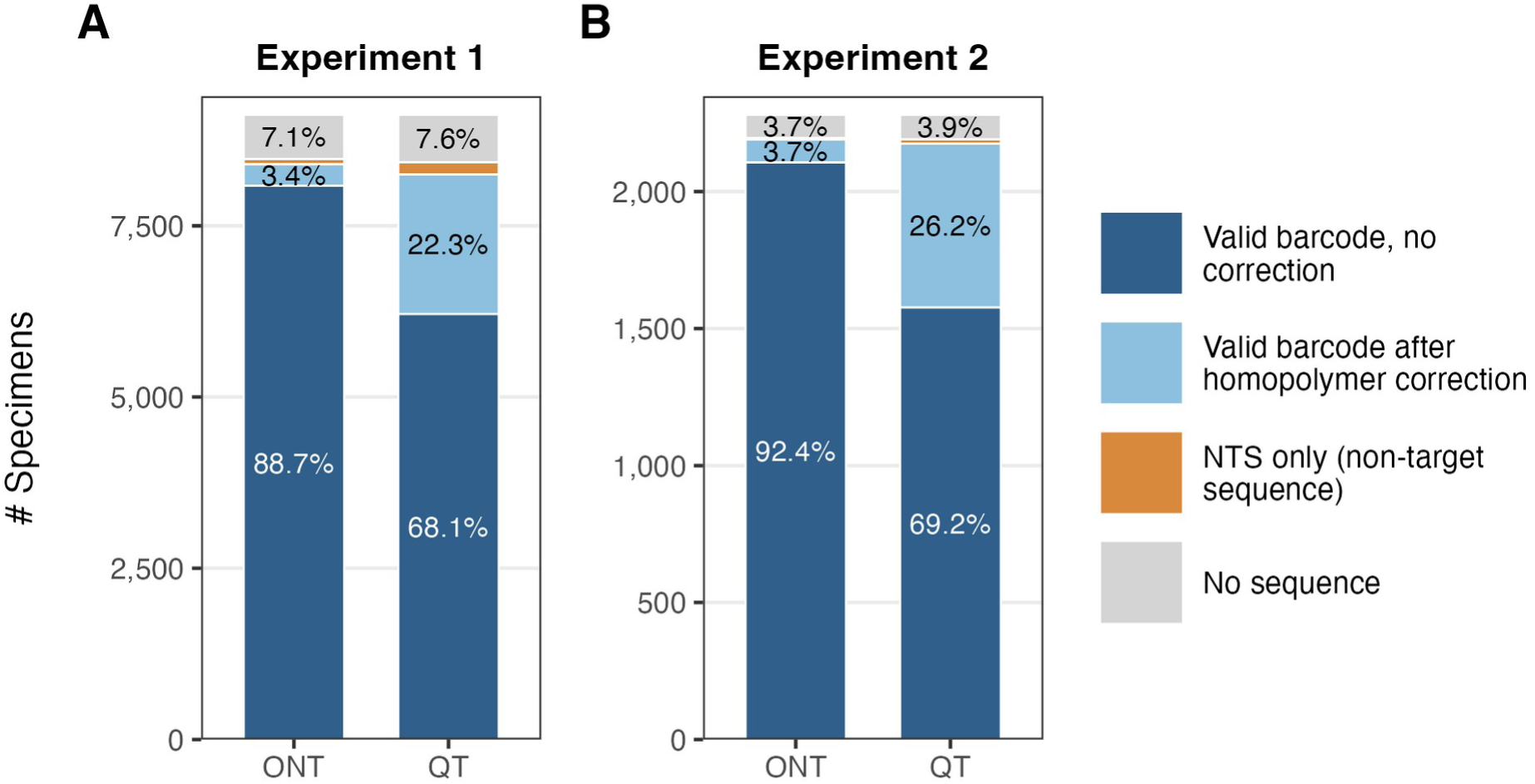
Barcode outcome for every specimen in (A) Experiment 1 (n = 9,120) and (B) Experiment 2 (n = 2,280), by platform. Valid barcodes are split by whether BIP’s homopolymer error correction edited the sequence (adding Ns, or removing extra bases). “NTS only (non-target sequence)” marks specimens with no valid barcode where an OTU matching the expected order failed only because of a frameshift indel that was not corrected (the well’s only sequences were non-target sequences, NTS); “No sequence” covers all other specimens without a valid barcode. Negative controls are excluded; y-axes are scaled separately for each experiment.

**Table 1.** Experiment 1 Summary. Both ONT and QT runs analyzed a pool of 9,120 specimens and had 3.07M reads in the input file (mean Q ≥ 10; the ONT run was downsampled to match). “% retained” is the percent of input reads. Valid barcodes with ambiguous bases are given as a percent of valid barcodes.

|  | *ONT (MinION)* | *QT (QCell)* |
|---|---|---|
| *Reads in OTUs (% retained)* | 2,035,597 (66.3%) | 1,512,516 (49.3%) |
| *# Valid Barcodes (% success)* | 8,398 (92.1%) | 8,249 (90.4%) |
| *Valid barcodes with ambiguous bases* | 335 (4.0%) | 2,055 (24.9%) |
| *Mean read depth of barcodes (±SD)* | 222.3 (±97.2) | 163.1 (±69.0) |

### Experiment 2: Performance of QCell and Flongle flow cells

The partial QCell run generated 891,015 reads with a mean Q-score ≥ 10 and was downsampled to the 859,872 generated by the Flongle (Table S4; see Table S1 for non-downsampled analysis). BIN assignments were identical for 99.3% of specimens with a barcode on both platforms and a BIN on at least one (Table S7). ONT generated valid barcodes for 2,190 (96.1%) specimens while QT generated valid barcodes for 2,175 (95.4%) (Table 2; Figure 2B). With all 891,015 reads, the QCell recovered 2,176 barcodes (95.4%), the same as at equal depth (Table S1). We attribute the higher mean barcode recovery (e.g., ONT: 96.1% vs. 92.1%) compared to Experiment 1 to the different template DNA used. Diagnostics of the recovered sequences, and patterns of BIN and taxonomic assignment, were similar to what we observed in Experiment 1 (Fig. S5; Tables S6–S7).

**Table 2.** Experiment 2 Summary. Both ONT and QT runs analyzed a pool of 2,280 specimens and had 859,872 reads (mean Q ≥ 10) in the input file; the QCell run (891,015 reads) was downsampled to match the Flongle run. “% retained” is the percent of input reads. Valid barcodes with ambiguous bases are given as a percent of valid barcodes.

|  | *ONT (Flongle)* | *QT (QCell)* |
|---|---|---|
| *Reads in OTUs (% retained)* | 519,977 (60.5%) | 486,378 (56.6%) |
| *# Valid Barcodes (% success)* | 2,190 (96.1%) | 2,175 (95.4%) |
| *Valid barcodes with ambiguous bases* | 88 (4.0%) | 596 (27.4%) |
| *Mean read depth of barcodes (±SD)* | 225.9 (±84.4) | 211.1 (±76.4) |

### Experiment 3: Effectiveness of QCell flow cell wash

Each run generated just over 100K reads before it was terminated by QPreasy. Initial pore count declined from 223 to 112 across the eight runs, leading, as expected, to an increase (24 to 53.5 min) in the time required to achieve 100K reads (Table 3; Fig. S4C–D). The initial decline was slight, but it hastened after four runs. We detected four reads (two from each of two OTUs) in Run 4 that originated in Run 3 (Table 3), suggesting very infrequent carryover. We also detected one OTU with two reads in Run 3 that was misassigned to a Run 5 well, most likely through index hopping (Table 3). Because two reads per OTU is below the minimum recommended read count for nanopore data, there would be no detectable carryover under these parameters. This experiment established that washing the QCell prevented meaningful carryover as each run recovered the expected sequence in nearly all wells.

**Table 3.** Experiment 3 Summary. The same flow cell was used for all eight runs. Each run was programmed to end at 100K reads, and 100K reads were analyzed for each. “OTUs from other plates” counts OTUs (reads in parentheses) assigned to wells of any plate other than the one sequenced in that run.

| *Run #* | *Active pores at start* | *Time to 100K reads (min)* | *Reads in final OTUs* | *Barcodes recovered* | *OTUs from other plates (reads)* |
|---|---|---|---|---|---|
| *1* | 223 | 24 | 54,361 | 96 | 0 (0) |
| *2* | 219 | 26 | 54,454 | 96 | 0 (0) |
| *3* | 217 | 27 | 57,216 | 95 | 1 (2)<sup>†</sup> |
| *\*4* | 198 | 31.5 | 53,054 | 94 | 2 (4)<sup>†</sup> |
| *5* | 199 | 34.5 | 48,622 | 95 | 0 (0) |
| *6* | 163 | 40.5 | 48,415 | 96 | 0 (0) |
| *7* | 153 | 43 | 49,740 | 92 | 0 (0) |
| *8* | 112 | 53.5 | 51,140 | 96 | 0 (0) |

\*Runs 4–8 were conducted 23 days after runs 1–3.
<sup>†</sup>OTUs assigned to wells of another run’s plate. Run 3: 1 OTU (2 reads) of *Hermetia illucens*, assigned to a well of the Banded Cricket plate (Run 5). **The species does not match the plate (it is the species sequenced in Run 3 itself), so this is not carryover; it is probably index hopping or a similar misassignment.** Run 4: 2 OTUs (4 reads) of *Hermetia illucens*, assigned to wells of the Black Soldier Fly plate (Run 3). **The species matches the plate, consistent with carryover from Run 3.**

### Experiment 4: Ultra-fast library preparation

The ultra-fast kit enabled library preparation in a single step. Table 4 shows that a higher proportion (74.7% vs. 43.4%; 1.7×) of the reads from the ultra-fast kit were filtered during bioinformatic processing than those from the standard kit. However, the impact on barcode recovery was modest as the standard kit generated barcodes from 2,175 (95.4%) specimens while the ultra-fast kit generated them from 2,126 (93.2%) (Table 4; Table S2). Using all 3.85M reads from the run (4.5× the equal-depth input) raised recovery to 2,173 (95.3%), almost matching the standard kit with all of its reads (2,176; 95.4%; Table S2).

**Table 4.** Experiment 4 Summary. The left column is repeated from Table 2 to facilitate comparison. Both runs had 859,872 reads in the input file (mean Q ≥ 10). “% retained” is the percent of input reads. Valid barcodes with ambiguous bases are given as a percent of valid barcodes.

|  | *QT (Standard [Expt 2])* | *QT (Ultra-Fast)* |
|---|---|---|
| *Reads in OTUs (% retained)* | 486,378 (56.6%) | 217,133 (25.3%) |
| *# Valid Barcodes (% success)* | 2,175 (95.4%) | 2,126 (93.2%) |
| *Valid barcodes with ambiguous bases* | 596 (27.4%) | 655 (30.8%) |
| *Mean read depth of barcodes (±SD)* | 211.1 (±76.4) | 90.7 (±38.2) |

### Comparison of QT and ONT data quality

In Experiments 1 and 2, ONT generated barcodes for 10,588 specimens and QT generated barcodes for 10,424—98.5% as many as ONT (Tables 1–2). Without BIP’s homopolymer sequence correction, QT’s success would have been far lower (23.7% and 27.1% of QT barcodes were rescued by correction) (Figure 2; Table S8). We now compare data quality between ONT and QT to examine the reason for this small difference. Specifically, we examined: (i) the impact of local variation in GC composition on the incidence of frameshift indels for both ONT and QT; and (ii) overall read accuracy.

The consensus sequences generated from QT data consistently had more ambiguous bases than those from ONT. These ambiguous bases were added during bioinformatic processing to correct 1–2 bp frameshift indels in homopolymer regions (Fig. S6; Table S9). Moreover, the difference was markedly elevated in high-GC windows—rising to 0.22% (95% CI 0.21–0.23%), about 1 in 450 positions, in windows of ≥83% GC (ONT: 0.003%) (Figure 3A; Table S10). By contrast, ONT produced the highest incidence of ambiguous bases in low-GC regions, but its error rate was still less than for QT. Nearly all Ns in QT barcodes (96.9% in Experiment 1 and 99.0% in Experiment 2) were in or next to a homopolymer, although such positions made up only about 16% of the barcode (Table S9). Most QT errors were deletions masked by Ns, concentrated in C and G homopolymers of five or more bases (Fig. S6). These Ns rarely affected identification, as QT barcodes containing Ns were assigned the same BIN as ONT in 99.5% (Experiment 1) and 99.8% (Experiment 2) of cases (Table S9).

**Figure 3.**
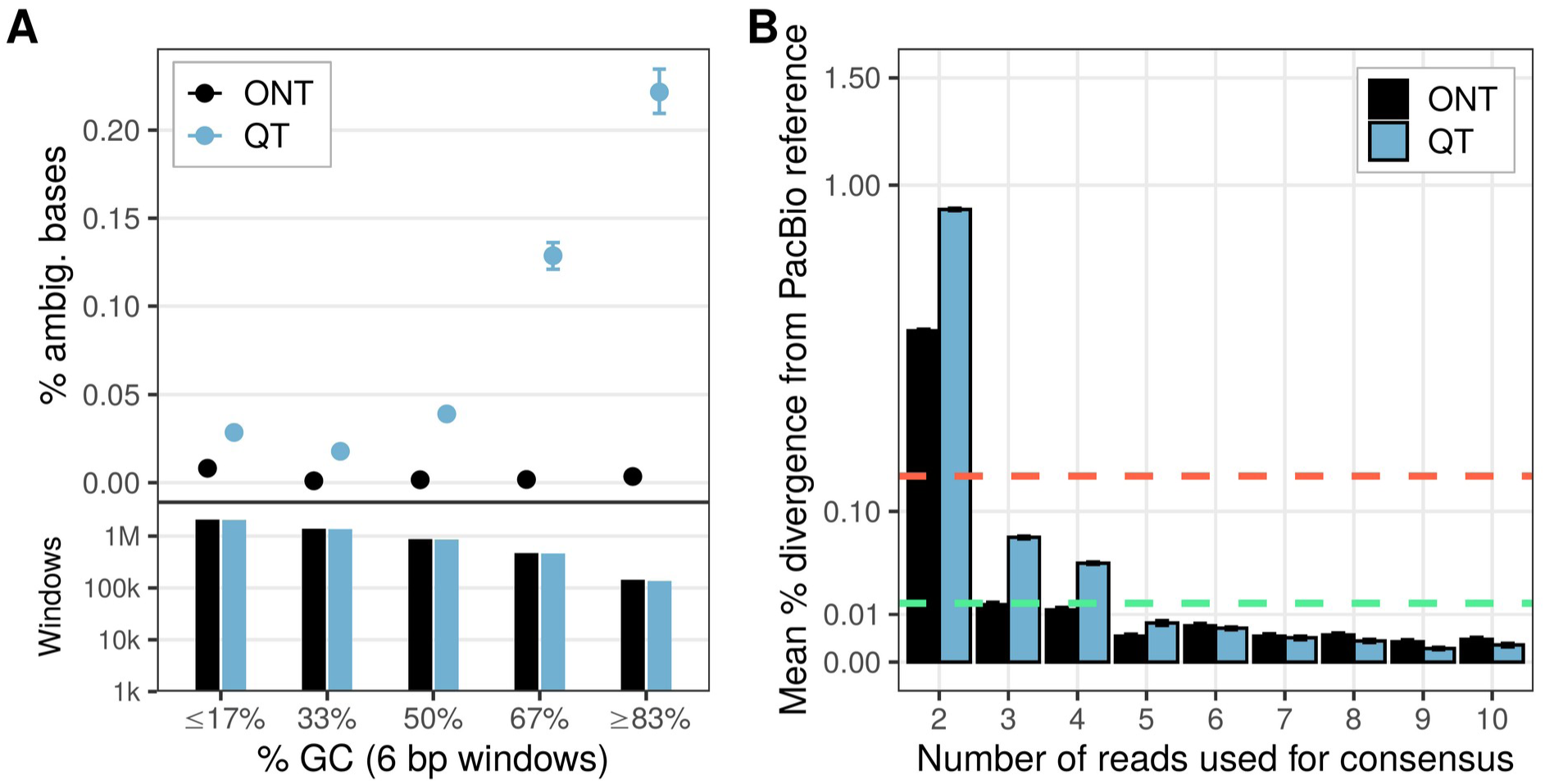
Ambiguous base calls and read accuracy in Experiment 1 for ONT (MinION; black) and Qitan (QCell; blue). (A) Incidence of ambiguous base calls in 6 bp windows with varying GC composition, from valid barcodes of length 658 (ONT n = 7,649; QT n = 7,515). Every overlapping 6 bp window of each barcode (653 per barcode) was assigned to a GC bin by its number of G or C bases (an N counts as neither). Points give the % ambiguous bases in each bin: N bases in that bin’s windows divided by all bases in those windows. Error bars are 95% confidence intervals from 2,000 bootstrap resamples of barcodes within platform (barcodes rather than windows were resampled because overlapping windows from one barcode are not independent). The lower panel gives the number of windows in each bin (log scale): ≤ 17%, 2,098,913 (ONT) and 2,064,932 (QT); 33%, 1,399,812 (ONT) and 1,375,535 (QT); 50%, 879,429 (ONT) and 863,948 (QT); 67%, 472,574 (ONT) and 465,617 (QT); ≥ 83%, 144,069 (ONT) and 137,263 (QT); in total, 4,994,797 (ONT) and 4,907,295 (QT) windows were analysed. Both ends of the GC range are pooled (0–1 and 5–6 G/C of 6 bases): the 100% bin is small and cannot contain an N (see below), and the low end is pooled in the same way for symmetry. Taken alone, the 100% GC windows (ONT 16,434; QT 13,911) contain no ambiguous bases. This is expected rather than a lower error rate: BIP inserts an N where a base is missing at the end of a homopolymer run (75% of QT Ns lie directly beside a run of three or more G or C bases), and a window containing an N has at most five G or C bases, so it is counted in the 83% rather than the 100% bin. Per-bin values, including the single-count bins, are in Table S10. (B) Evaluation of read accuracy. Analysis included all 5,399 specimens where ONT, QT and the PacBio SMRT barcode in BOLD (dataset DS-QT96) were the same 658 bp sequence with no ambiguous bases, with at least 10 reads supporting it on each of ONT and QT. The mean % sequence distance from the PacBio reference is shown on the y-axis for varying numbers of underlying sequences. The y-axis is on a square-root scale. The red dashed line reflects one error in a 658 bp sequence. The green dashed line reflects one error in ten 658 bp sequences. Error bars represent ±1 SE.

Analysis of individual component reads revealed that sequences generated by QT were slightly less accurate than those from ONT (Figure 3B; Fig. S3). While ONT data more closely resembled the PacBio reference for read depths of 2–4, both sequencers delivered a mean of less than 1 error per sequence when the consensus was based on three reads. Both ONT and QT achieved fewer than 1 error per 10 sequences (0.1 errors per sequence), ONT with four reads and QT with five (Figure 3B). In Experiment 2, the Flongle and QCell performed similarly at every read depth, both reaching these thresholds with three and five reads, respectively (Fig. S3). We analyzed the per-read accuracy of both platforms and found that the MinION had the highest accuracy at 99.39%, followed by Flongle at 98.79% followed by QCell at 98.34% for Experiment 2 and 98.04% for Experiment 1 (median per-read identity; Tables S11–S12; Figs. S7–S8). QT’s lower per-read accuracy in Experiment 1 reflected an excess of deletions (0.88 vs. 0.26 per 100 aligned bases) and insertions (0.55 vs. 0.16), whereas substitution rates were closer (0.60 vs. 0.42; Table S11; Fig. S7). In Experiment 2, the Flongle had more substitutions than the QCell (0.63 vs. 0.54) but fewer insertions and deletions (Table S12; Fig. S8).

## DISCUSSION

This study has shown that the QNome sequencer paired with a QCell enables reliable DNA barcode recovery. As such, QT provides an effective alternative to ONT. In fact, the QNome has gained adoption by the CBG as its primary platform for experiments requiring <200K reads as the cost per run is well below that achievable with the Flongle when QCells are reused (about US$35 vs. US$210 per run; Table S5). Once Flongle is discontinued, the QCell will represent the only low-throughput (<10 Gb) nanopore flow cell well-suited for small-scale barcoding studies.

Overall read accuracy was lower for QT than ONT and this difference was particularly evident in regions with a GC content >50%. This difference may constrain the use of QT in some contexts, and it affected our results even though arthropod mitogenomes have low GC content (Cameron, 2014). Specifically, the mean GC content of the taxa examined in our study averaged 30.9% with just 1.2% of specimens producing a COI amplicon with ≥ 40% GC (Table S13, Fig. S9). Both platforms required an average of three reads to produce sequences with fewer than one error per sequence and five (QT) or four (ONT) reads to produce sequences with an average of less than one error per ten sequences (Figure 3B). Thus, minimum read depth required to achieve high-fidelity sequences does not differ significantly between the platforms. Nevertheless, QT’s relatively poor performance in GC rich regions is clearly the biggest barrier that must be overcome and is primarily responsible for the lower success rate than ONT. A significant fraction of specimens—approximately one-in-four—would not have had valid barcodes without our bespoke error correction (Table S8). Our success with QT might not apply to other taxa and markers with higher mean GC content, so further study is warranted.

BIP’s approach to homopolymer error correction was originally developed for ONT data. Examining our data, we found that a much larger fraction of specimens were ‘rescued’ by error correction for QT data than ONT data (Figure 2; Table S8). Thus, we recommend analysis of QT data for COI amplicon sequencing leverage our error correction approach. Further refinement of the error correction script— which is beyond the scope of this study—could further improve target barcode assignment. QT continues to improve the accuracy of its flow cells (pers. comm.) which could close the gap with ONT.

QT’s use for nuclear genome sequencing might prove more challenging. In their study of roughly 750 nuclear genomes, Frey & Pucker (2020) found that average GC content ranged from 47.1% in fungi to 39.4% in animals and 36.3% in plants. Although animal mitogenomes are a favourable setting for the use of QT, its performance should now be tested with other taxa and markers.

Our comparison of the two QT protocols for library preparation revealed a trade-off between speed and technical ease versus data quality and cost. Library preparation is, by far, the most challenging step in the nanopore sequencing workflow (Floyd et al., 2023). Because the ultra-fast protocol is much simpler to execute, it greatly reduces the risk of technical errors. The ultra-fast kit is more expensive (1.75×; QT Pers. Comm.) than the standard kit, but as it simplifies and speeds library preparation, it is an attractive option in time-limited settings such as field courses, especially when personnel with molecular expertise are unavailable. Sequence quality is lower, but this can be countered by generating more reads (Table S2).

The ability to reuse QCells after washing is a key advantage for small and/or iterative sequencing runs. The MinION flow cell can also be washed (Meyer et al., 2026) for the same cost as QCell, but the MinION protocol takes 60 min vs 20 min for QT. If the user wishes to acquire a target number of reads, multiple runs can be rapidly completed with a QCell. Because it requires 24–54 minutes to generate 100K reads (Table 3), six runs can be completed in eight hours. Our results indicate that washing degrades the flow cell gradually, as active pores declined from 223 to 112 over eight runs while output per pore remained stable (Table 3; Fig. S4C–D). Moreover, we found that the QCell could be set aside for at least three weeks between reusage without damaging its performance (Fig. S4D). This flexibility enables users to tailor sequencing effort to project size, reducing per-sample costs.

Our study focused only on COI-based DNA barcoding of arthropod specimens. It would be beneficial for future studies to compare QT and ONT performance via features not considered here. While exceeding the capacity of some short-read sequencers (Elbrecht et al., 2019), our amplicons do not test the performance of ONT (Jain et al., 2016) and QT (Song et al., 2025) with longer amplicons. Our study also does not explore native (i.e., PCR-free) DNA or RNA sequencing nor any epigenetic modification (Jenjaroenpun et al., 2021; Liu & Conesa, 2025). There is, clearly, much more comparative work awaiting QT’s devices.

Although its read quality is lower than that of the latest MinION flow cell (ONT R10.4.1), QT is an effective platform for DNA barcoding. It currently achieves read accuracies far exceeding those generated by ONT when ONT was first adopted for barcoding (Delahaye & Nicolas, 2021; Pomerantz et al., 2018; Silvestre-Ryan & Holmes, 2021) and it is possible that software and/or hardware updates will enhance its performance. Given ONT’s withdrawal of the Flongle, the high cost of the MinION flow cell and associated chemistry, and the risk of further retreat from affordable sequencing, QT’s entry into the nanopore sequencing market is a positive development. Once its products are widely available, QT is well positioned to support the realization of a distributed network of facilities employing DNA-based analysis to advance biodiversity science.

## AUTHOR CONTRIBUTIONS

- RF: Lead laboratory work

- SP: Bioinformatics; Laboratory work

- EO: Bioinformatics

- KT: Data analysis; Lead manuscript writing; Coordinate project

- SJ: Assist laboratory work

- PH: Project initiation and supervision; Funding acquisition; Lead manuscript revision

- All authors: Manuscript revisions

## ACKNOWLEDGEMENTS

We thank Ethan Duffy for managing the financial particulars of purchasing the sequencer and consumables from QT. We thank Jaclyn McKeowan and Suz Bateson for their aid in preparing Figure 1. We are very grateful to CBG staff for DNA extraction and PCR. Comments from Rodrigo Nunes-da-Fonseca, Lara Urban, and an anonymous reviewer improved the manuscript. This study was enabled by support to PDNH from the Government of Canada through Genome Canada and Ontario Genomics (OGI-208 and OGI-233), and by the New Frontiers in Research Fund (NFRFT-2020-00073). Infrastructure required to carry out the work was acquired with support from the Canada Foundation for Innovation (CFI) and the Gordon and Betty Moore Foundation. A Major Science Infrastructure award (MSI 42450) from CFI plays a key role in sustaining the CBG’s informatics and sequencing platforms.

## DATA ACCESSIBILITY

Raw nanopore reads from all sequencing runs are deposited in the NCBI Sequence Read Archive under BioProject PRJNA1561528. Specimen records and PacBio reference barcodes are in BOLD datasets DS-QT96 and DS-QT24. BIP v1.1.0 is available at https://github.com/cbg-innov/BIP. The complete reproducibility package (all data except raw reads, including BIP parameter files, analysis scripts, run reports and logs, and runs.tsv, which records the random seeds used for subsampling) is archived at Zenodo (https://doi.org/10.5281/zenodo.24204998).

## USE OF ARTIFICIAL INTELLIGENCE

Artificial Intelligence was used in the following ways:

- Proofreading manuscript text
- Producing analysis code and technical documentation
- Facilitating data archiving

We used Claude desktop App v2.42304.2 and mobile App v1.261016.20 with models Sonnet 5.5 and Opus 5.5. AI is a tool used by the authors who remain responsible and accountable for all content. All AI-assisted code was reviewed and tested by the authors, and AI was not used to interpret results or draft manuscript content.

## Supplementary Text S1. Qitan Tech (QT) Molecular Protocols

Because QT protocols are not currently publicly available, we summarise them here. In all cases, we followed the manufacturers protocols exactly.

### 1. DNA Library Prep Kit (QDL-E V3.0)

Product No. O-0041. 6 rxns per kit. Manual version 1.0.

Ligation-based library prep for genomic DNA, fragmented DNA or PCR amplicons. End repair, SAC ligation, bead cleanup, elution, quantification.

#### 1.1 Kit components

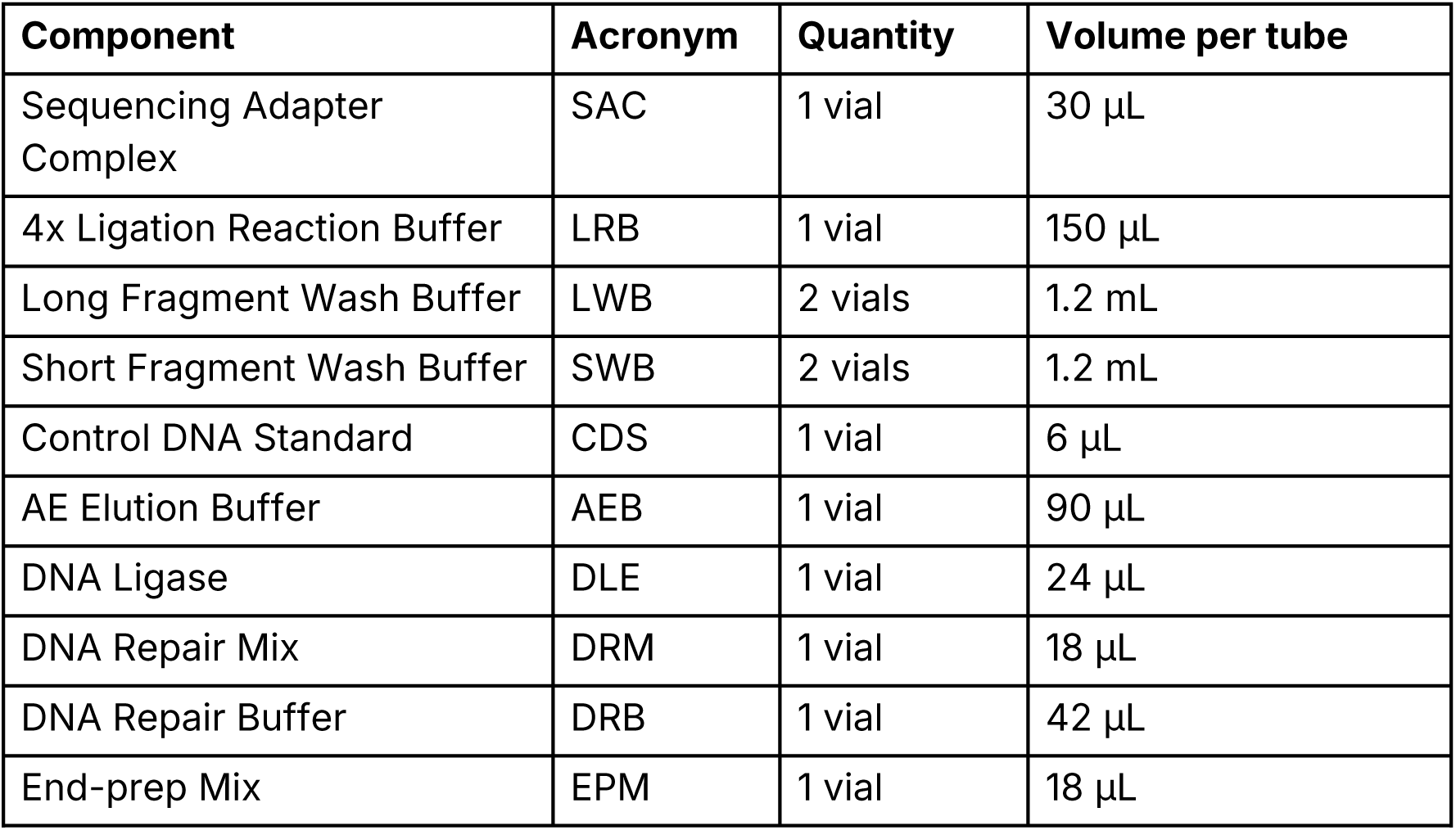

Store at -25 to -15 °C. Limit to 6 freeze-thaw cycles.

#### 1.2 User-supplied materials

Thermal cycler, magnetic rack (PCR and 1.5 mL tubes), mini centrifuge, vortex mixer, pipettes, ice, Qubit fluorometer. Anhydrous ethanol, nuclease-free water, VAHTS DNA Clean Beads or equivalent, Qubit dsDNA HS assay kit. PCR tubes and 1.5 mL tubes.

#### 1.3 Input

- Genomic DNA: 1 µg, roughly 150 to 300 fmol by Qubit. A260/A280 1.8 to 2.0, A260/A230 2.0 to 2.2. Fragmenting with a g-Tube first improves yield.
- Amplicon dsDNA: 150 to 300 fmol.

#### 1.4 Preparation

1. Flick to mix EPM, DRM, DLE and SAC individually, spin down, hold on ice.
2. Thaw the other reagents at room temperature, vortex 10 s, spin down, hold on ice.
3. Equilibrate the beads at room temperature for at least 30 min.
4. Prepare fresh 80% ethanol.

#### 1.5 DNA repair and end repair

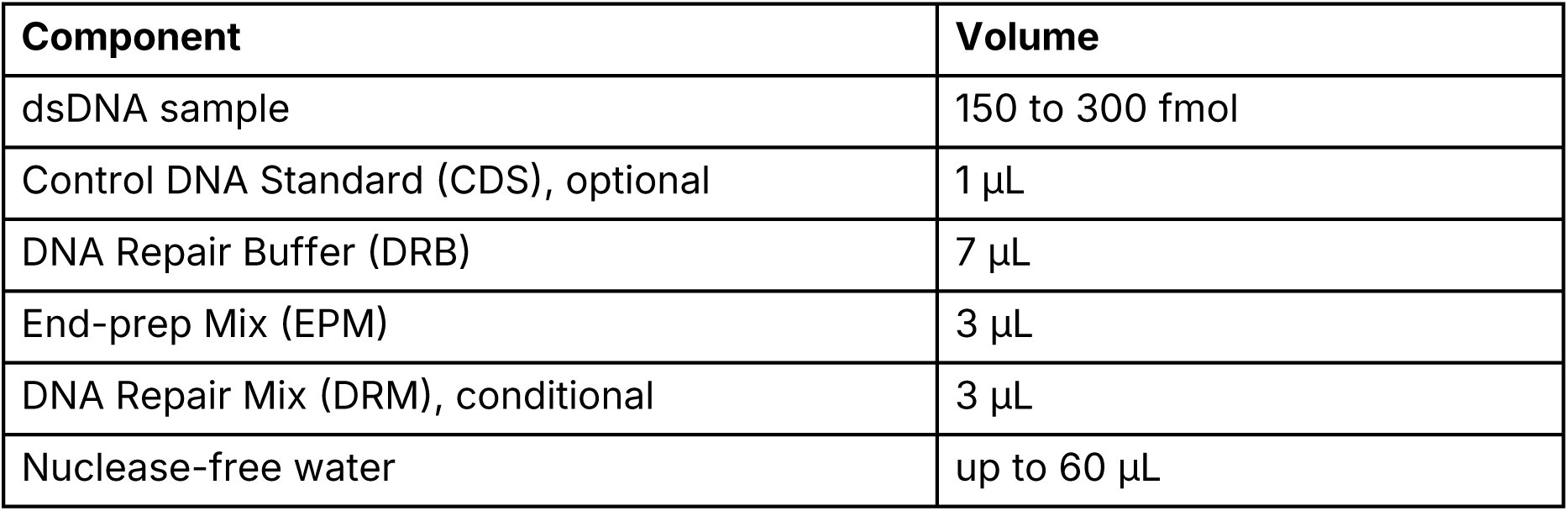

CDS: add 1 µL only when monitoring library quality. Pass criterion is CDS accuracy above 95%.

DRM: include for mechanically fragmented input, omit for PCR amplicons.

Flick to mix, spin down, then run: 30 °C for 10 min (20 °C if DRM omitted), 65 °C for 10 min with the lid at 75 °C, hold at 12 °C.

#### 1.6 Cleanup of end-repair products

1. Vortex the room-temperature beads for 30 s.
2. Add 60 µL beads to the 60 µL reaction (1:1), flick to mix, 5 min at room temperature.
3. Spin down, place on the magnet, and once clear (2 to 3 min) discard the supernatant with the tube on the magnet.
4. On the magnet, add 200 µL fresh 80% ethanol without disturbing the pellet, wait 30 s, discard.
5. Repeat step 4 once.
6. Spin down, return to the magnet, remove all residual liquid, air dry with the lid open until the beads are matte (3 to 5 min).
7. Off the magnet, resuspend in 60 µL nuclease-free water, 3 min at room temperature.
8. Spin down, return to the magnet, and once clear transfer the supernatant to a fresh 1.5 mL tube.

#### 1.7 Adapter ligation

| **Component** | **Volume** |
|---|---|
| Purified end-repair product | 60 µL |
| Sequencing Adapter Complex (SAC) | 5 µL |
| 4x Ligation Reaction Buffer (LRB) | 25 µL |
| DNA Ligase (DLE) | 4 µL |
| Nuclease-free water | up to 100 µL |

Flick to mix, spin down, 10 min at room temperature. Do not vortex.

#### 1.8 Cleanup of ligation products

1. Add beads: 100 µL (1:1) for fragments 200 bp or shorter, 40 µL (1:0.4) for longer than 200 bp. Flick to mix, 5 min at room temperature.
2. Spin down, pellet on the magnet until clear and colourless, pipette off the supernatant.
3. Off the magnet, resuspend in 200 µL wash buffer: LWB for fragments longer than 3 kb, SWB for shorter than 3 kb. Flick, spin down, return to the magnet, remove the supernatant once pelleted.
4. Repeat step 3 once.
5. Spin down, return to the magnet, remove residual liquid. Off the magnet, resuspend in 15 µL AEB, 5 min at room temperature.
6. Spin down, pellet on the magnet until clear, transfer the eluate (the library) to a clean 1.5 mL tube.

#### 1.9 Quantification

Qubit dsDNA HS assay. Convert mass concentration to molar before loading.

#### 1.10 Kit-specific notes

- Remove all residual ethanol after the end-repair cleanup or ligation will be compromised.
- Mix by flicking during ligation and when pooling libraries. Do not vortex.
- Take beads out of the fridge at least 30 min before use.
- Resuspend the beads fully during the ethanol washes.

### 2. DNA Sequencing Kit (QDS V3.0)

Product No. O-0050. 6 rxns per kit. Manual version 2.1.

Primes the flow cell and loads the library for sequencing.

#### 2.1 Kit components

| **Component** | **Acronym** | **Quantity** | **Volume per tube** |
|---|---|---|---|
| Sequencing Reaction Buffer | SRB | 1 vial | 540 µL |
| Priming Buffer | PRB | 3 vials | 1.2 mL |
| Priming Anchor | PRA | 1 vial | 24 µL |

Store at -25 to -15 °C.

#### 2.2 User-supplied materials

QNome-3841 or QNome-3841hex, or QPursue-6k or QPursue-6khex sequencer. Mini centrifuge, pipettes, vortex mixer. Nuclease-free water. QCell V3.0 flow cell, 1.5 mL tubes.

#### 2.3 Flow cell priming

1. Thaw PRA, SRB and one tube of PRB completely at room temperature.
2. Vortex and invert PRA, SRB and PRB to mix.
3. Peel off the sealing film and turn the rotary valve clockwise about 90 degrees until it stops.
4. Set a 200 µL pipette to 185 to 190 µL, insert the tip into the loading port, turn the knob roller counterclockwise to draw out air bubbles and a small volume of liquid.
5. Draw 200 µL PRB, expel any bubble at the tip, insert into the loading port, dispense slowly and evenly until 2 to 5 µL remains in the tip.
6. Repeat step 5 once.
7. Combine 4 µL PRA with 200 µL PRB in an Eppendorf tube, vortex, spin down.
8. Draw 200 µL of the PRA/PRB mix, expel any bubble, load slowly until 2 to 5 µL remains in the tip. Avoid air bubbles.
9. Turn the rotary valve clockwise about 90 degrees until it stops, reapply the sealing film, stand 5 min (QCell-384 V3.0) or 15 min (QCell-6k V3.0).

#### 2.4 Library loading mix, QNome platform

| **Component** | **Volume** |
|---|---|
| Sequencing Reaction Buffer (SRB) | 35 µL |
| DNA library | V µL |
| Nuclease-free water | up to 70 µL |

- Under 1 kbp: 20 to 30 fmol
- 1 to 20 kbp: 15 fmol
- Over 20 kbp: 100 to 200 ng

#### 2.5 Library loading mix, QPursue platform

| **Component** | **Volume** |
|---|---|
| Sequencing Reaction Buffer (SRB) | 90 µL |
| DNA library | V µL |
| Nuclease-free water | up to 180 µL |

- Under 1 kbp: 60 to 80 fmol
- 1 to 2 kbp: 40 to 60 fmol
- 2 to 20 kbp: 35 fmol
- Over 20 kbp: 100 to 200 ng

#### 2.6 Loading volume

V = n x 0.62 x N / C

V = volume of library to load in µL, n = loading amount in fmol, N = fragment length in kbp, C = library concentration in ng/µL.

For libraries built with the Ultra-fast kit (QSP), use the loading amounts in Section 4.6 instead of the tables above.

#### 2.7 Loading and run start

1. Peel off the sealing film from the loading port and turn the rotary valve clockwise about 90 degrees until it stops. Add the loading mix slowly and evenly until 2 to 5 µL remains in the tip.
2. Turn the rotary valve counterclockwise about 90 degrees until it stops and reapply the sealing film.
3. Set the waiting time to 0 min in the dialog box and click Confirm to start sequencing. For short-fragment samples with a short run time, use 5 to 15 min instead.

### 3. Flow Cell Wash Kit (QNW V3.2)

Product No. O-0048 (6 rxns) and O-0049 (48 rxns). Manual version 1.0.

Digests and flushes the loaded library so a flow cell can be reused or stored. Compatible with QCell-384 and Qcell-6k.

#### 3.1 Kit components, 6 rxns per kit

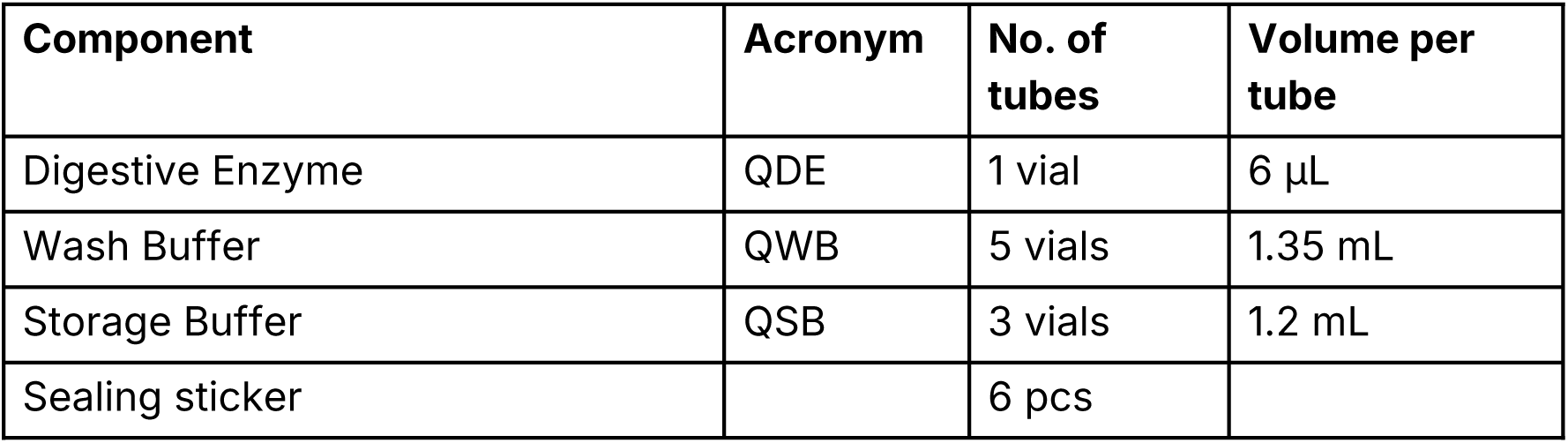

#### 3.2 Kit components, 48 rxns per kit

| **Component** | **Acronym** | **No. of tubes** | **Volume per tube** |
|---|---|---|---|
| Digestive Enzyme | QDE | 1 vial | 48 µL |
| Wash Buffer | QWB | 1 tube | 53 mL |
| Storage Buffer | QSB | 1 tube | 29 mL |
| Sealing sticker |  | 48 pcs |  |

Store at -25 to -15 °C. Reseal promptly after opening.

### 3.3 User-supplied materials

Mini centrifuge, vortex mixer, pipettes, timer, ice. 1.5 mL Eppendorf tubes.

#### 3.4 Reagent setup

1. Flick QDE to mix, spin down, hold on ice.
2. Thaw QWB and QSB completely at room temperature, vortex 10 s, spin down, hold at room temperature.
3. Close the rotary valve of the flow cell (vertical position). Insert a 200 µL pipette into the waste port, draw off all waste buffer, discard.
4. Combine 1 µL QDE with 99 µL QWB in a 1.5 mL tube, flick to mix, spin down.
5. In a new 1.5 mL tube, dilute the step 4 mixture by fragment length, then flick to mix and spin down:

| **Component** | **1 kb or shorter** | **Longer than 1 kb** |
|---|---|---|
| Mixture from step 4 | 20 µL | 4 µL |
| Wash Buffer (QWB) | 980 µL | 996 µL |

Prepare the step 4 and step 5 mixtures fresh each time.

#### 3.5 Washing, QNome platform

a. Peel back part of the sealing sticker and turn the rotary valve clockwise about 90 degrees until it stops.
b. Load 200 µL of the step 5 mixture through the loading port, pushing the plunger down slowly and leaving a small volume in the tip to avoid air bubbles.
c. Repeat step b twice, three loads in total.
d. Close the rotary valve (counterclockwise), reapply the sticker, stand 10 to 20 min at room temperature.

#### 3.6 Washing, Qpursue platform

a. Peel back part of the sealing sticker and turn the rotary valve clockwise about 90 degrees until it stops.
b. Load 200 µL of the step 5 mixture as above, leaving a small volume in the tip.
c. Repeat step b three times, four loads in total.
d. Close the rotary valve (counterclockwise), reapply the sticker, stand 20 min at room temperature.

#### 3.7 Inactivation, Qnome platform

a. Peel back part of the sealing sticker and turn the rotary valve clockwise about 90 degrees until it stops.
b. Load 200 µL QSB through the loading port, leaving a small volume in the tip.
c. Repeat step b once, two loads in total.
d. Close the rotary valve (counterclockwise), reapply the sticker, stand 20 min at room temperature.

#### 3.8 Inactivation, Qpursue platform

a. Peel back part of the sealing sticker and turn the rotary valve clockwise about 90 degrees until it stops.
b. Load 200 µL QSB as above, leaving a small volume in the tip.
c. Repeat step b twice, three loads in total.
d. Close the rotary valve (counterclockwise), reapply the sticker, stand 20 min at room temperature.

#### 3.9 Finishing

1. Confirm the rotary valve is closed. Insert a 200 µL pipette into the waste port, draw off all waste buffer, discard.
2. To store the flow cell, place it at 4 to 8 °C. The 20 min room-temperature stand in Sections 3.7 and 3.8 is then not required. To sequence a new library immediately, continue from Section 2.

#### 3.10 Kit-specific notes

- QWB and QSB must be at room temperature before use.
- Replace the sealing sticker if it loses adhesion.

### 4. Ultra-fast DNA Library Prep Kit (QSP V3.0)

Product No. O-0042. 6 rxns per kit. Manual version 1.2.

Rapid amplicon library prep, roughly 9 min. Amplicons are A-tailed and ligated directly to the Rapid Enzyme Adapter with no cleanup in between. Samples barcoded during PCR can be multiplexed.

#### 4.1 Kit components

| **Component** | **Acronym** | **Quantity** | **Volume per vial** |
|---|---|---|---|
| A-Tailing Buffer | AB | 1 vial | 12 µL |
| A-Tailing Reagent 1 | A1 | 1 vial | 12 µL |
| A-Tailing Reagent 2 | A2 | 1 vial | 12 µL |
| Rapid Enzyme Adapter | REA | 1 vial | 60 µL |
| Washing Buffer | WB | 1 vial | 600 µL |
| Nuclease-Free Water | NFW | 1 vial | 84 µL |

Store at -25 to -15 °C.

#### 4.2 Input

PCR amplicons, isothermal amplification products or polymerase extension products. The 5′ terminal hydroxyl must be unmodified and the 3′ end blunt or A-overhang. A260/A280 1.8 to 2.0, A260/A230 2.0 to 2.2.

- QNome platform: 200 fmol total input
- QPursue platform: 600 fmol total input
- Molar conversion: fmol/µL = 1613 x ng/µL / median fragment length in bp.

#### 4.3 Preparation and bead washing

1. Invert and mix A1 and REA thoroughly, spin down, hold on ice. Thaw the other reagents at room temperature, shake 10 s, spin down, hold on ice.
2. Invert and gently flick REA until no beads remain aggregated. Transfer 10 µL REA per reaction to a tube, place on the magnet, and once clear discard the supernatant. Hold on ice.
3. Add 50 µL WB to the beads, mix by pipetting, place on the magnet, discard the supernatant once clear. Repeat twice. Cap, spin down, return to the magnet, remove any remaining WB, hold on ice.

#### 4.4 A-tailing reaction

| **Component** | **Volume** |
|---|---|
| Amplicons (200 fmol QNome, 600 fmol QPursue) | X µL |
| A-Tailing Buffer (AB) | 2 µL |
| A-Tailing Reagent 1 (A1) | 2 µL |
| A-Tailing Reagent 2 (A2) | 2 µL |
| Nuclease-free water (NFW) | up to 20 µL |

Flick to mix, spin down, incubate at room temperature for at least 20 s.

#### 4.5 Adapter ligation

1. Transfer the full 20 µL A-tailing reaction to the washed REA beads. Tap the tube wall gently to mix, incubate at room temperature for 5 to 20 min. More than 10 min is recommended. Tap periodically so the beads stay suspended.
2. Spin down, place on the magnet, and once clear transfer 19 µL of the supernatant (the library) to a fresh 1.5 mL tube.

#### 4.6 Sequencing

Follow Section 2 for priming, loading and run start, using these loading amounts in place of the fragment-length tables:

- QNome platform: 80 fmol, roughly 8 µL of library
- QPursue platform: 240 fmol, roughly 8 µL of library

#### 4.7 Kit-specific notes

- Mix the REA adapter and the library by gentle flicking only. Do not vortex.
- Flick gently during ligation to keep the beads from aggregating.

### 5. General precautions, all four kits

- Store all components at -25 to -15 °C and avoid repeated freeze-thaw cycles. Ship refrigerated at no warmer than -15 °C.
- Fully thaw and mix every component before use.
- Avoid cross-contamination between samples and products.
- Use filter tips and RNase-free and DNase-free tubes.
- Quantify by Qubit. Dilute samples above 100 ng/µL before measurement, with no single dilution step exceeding 100-fold. Inaccurate quantification degrades library quality.
- Flush any reagent spill with plenty of water.

## Supplementary Text S2. Barcode Inference Pipeline

### Installation and use

BIP is the standard workflow used by the Centre for Biodiversity Genomics for the analysis of DNA barcode data (Prosser et al., 2026). BIP is installed using Docker. This study used BIP v1.1.0. Running a container from the image file generates BIP’s working environment and installs specific versions of all required software. First- time users can execute a standardized command with trial data to confirm successful installation. To initiate analysis, BIP requires a single gzipped .fastq file (or a folder containing .fastq files), an Excel file (.xlsx) with parameters, and a reference library.

BIP’s parameters file allows the user to specify the specimen names, the UMIs, and the primers used for each run. In the parameters, users can also enter any collateral information they would like in the results files (e.g., geographic coordinates or collection information). Most collateral information is not used for analysis, except for taxonomic information. If the user enters taxonomy, this information is employed by BIP to distinguish target from non-target sequences. For instance, if the user specifies that a specimen belongs to the order Hemiptera but sequences are recovered from both a Hemiptera and a Pipunculidae (Diptera) parasitoid, the Hemiptera sequence is designated as the target while the Diptera is classed as the non-target.

BIP supports reference libraries as SINTAX-formatted .fasta files (R. Edgar, 2016). In the SINTAX format, the sample name is followed by taxonomic information in a standard format. Optionally, BINs can be included following a vertical bar (i.e., ‘|’) after the sample ID. For example, a valid header is: >Example_SampleID| BOLD:AAA0001;tax=k:Animalia,p:Chordata,c:Mammalia,o:Primates,f:Hominidae,g:Homo,s:Homo sapiens;

Any number of genetic markers and reference libraries can be used in a run, but this needs to be indicated in the input data.

### Workflow

For long-read sequencers (PacBio, ONT, QT), BIP accepts a single gzipped .fastq file as input, or a folder containing gzipped .fastq files. For paired-end (i.e., Illumina) data, a single R1 and R2 file must be provided. Data type is indicated as a command argument. BIP then orients reads, assigns them to primer pairs, filters out reads outside of the user-specified length interval and employs *cutadapt* (Martin, 2011) to demultiplex by UMIs. Next, any sequence (e.g., primers) flanking the target gene region is trimmed. Only those reads where both UMIs are found are retained.

Chimera removal follows filtering and trimming for high-quality data (PacBio and paired-end Illumina reads) and is accomplished using vsearch’s *de novo* chimera detection. Sequences are then trimmed once again to the size range expected for the target marker. For nanopore data (ONT and QT), chimera removal occurs after OTU clustering (see below).

Demultiplexed sequences are clustered into OTUs using vsearch’s clustering algorithm with a user-defined threshold (here 95%). Only those OTUs reaching a user-defined minimum read depth (here 5) are retained. A secondary clustering step with a user-defined threshold (typically higher; here 97%) is then used to refine the OTUs. Following secondary clustering, OTUs are filtered to the user’s specified minimum read depth. For each OTU, BIP then builds a consensus sequence from a homopolymer-aware MAFFT alignment of its distinct reads, weighted by read count (Katoh & Standley, 2013); very large OTUs are subsampled. Chimera removal occurs at this point for nanopore data.

After filtering by read depth, two COI-specific steps are invoked (only when the marker is specified as ‘COI-5P’). First, a custom error correction script is applied which involves aligning each sequence to its closest match in a reference database of sequences capturing all known amino acid sequences for COI (derived from BOLDistilled) (Prosser et al., 2025). Since sequencing errors occur primarily in homopolymer tracts, the error correcting script identifies any frameshift indels (1–2 bp) that are within homopolymers and adds ambiguous ‘N’ bases (or removes 1–2 extra bases) to restore the proper reading frame. Homopolymers are defined as four or more identical bases. Second, COI sequences are checked for stop codons. BIP does not attempt to correct gaps outside of homopolymers as those are more likely to reflect true biological variation than a sequencing error.Any sequence with a remaining frameshifting indel or stop codon is flagged as a NUMT and is designated as a non-target sequence.

After error correction and NUMT-flagging, taxonomy is assigned probabilistically using SINTAX (R. Edgar, 2016) in vsearch with a user-defined confidence threshold (here 0.6). If the marker is COI-5P (≥300 bp) and BINs are included in the SINTAX-formatted .fasta headers, BIP conducts BIN-matching. A match requires a user-defined identity (here 97%) over at least 75% of the amplicon, and BIP returns the closest matching BIN rather than the first acceptable hit. If an OTU does not match a BIN, its nearest-neighbor BIN is returned. The distance value for each OTU to its nearest neighbor is also returned.

OTUs are then designated as either ‘target’ or ‘non-target sequence’ (NTS). While a maximum of one target sequence is returned per specimen, there is no maximum number of NTS. An OTU must match the expected taxonomy and have no sequence issues (indels or stop codons) to be designated as a target. If multiple OTUs meet both criteria, the target is the OTU with the most specific taxonomic assignment, then the fewest ambiguous bases, then the length closest to that expected, and then the most reads. If two or more OTUs are still tied, then all are designated as NTS.

Results are returned to the user in a format that facilitates uploading to BOLD (if desired) in addition to various downstream analyses.

## Supplementary Text S3. DNA Extractions for Experiment 3

DNA extracts were obtained from the following species: Plate 1 DNA derived from larval *Galleria mellonella* (Pyralidae, Lepidoptera); Plate 2 from larval *Zophobas atratus* (Tenebrionidae, Coleoptera); Plate 3 from larval *Hermetia illucens* (Stratiomyidae, Diptera); Plate 4 from *Apis mellifera* (Apidae, Hymenoptera); Plate 5 from *Gryllodes sigillatus* (Gryllidae, Orthoptera); Plate 6 from *Drosophila melanogaster* (Drosophilidae, Diptera); Plate 7 from *Tribolium castaneum* (Tenebrionidae, Coleoptera); Plate 8 from *Blaberus discoidalis* (Blaberidae, Blattodea).

Bulk DNA was extracted from one specimen of each species using a standard silica membrane-based protocol (Ivanova et al., 2006) with modifications given below. Briefly, tissue was cut into small pieces using sterile scissors and placed in a 2 mL tube containing 1 mL of lysis buffer (700 mM guanidine thiocyanate, 30 mM EDTA pH 8.0, 30 mM Tris-HCl pH 8.0, 0.5% Triton X-100, 5% Tween-20, 2 mg/mL proteinase K). Samples were gently agitated at 56 °C for 2 h, after which 900 µL of each lysate was transferred into a 5 mL tube containing 1.8 mL of binding mix (Ivanova et al., 2006). DNA was then purified by applying 700 µL of the binding solution to a spin column (Epoch Biolabs) and centrifuging for 30 s. The flowthrough was discarded and the process was repeated until all 2.7 mL of binding mixture was passed through the column. The column membrane was then washed once with 500 µL of protein wash buffer (Ivanova et al., 2006) with the flowthrough discarded after each centrifugation. Residual wash buffer was removed via a final centrifugation followed by incubation at 56 °C for 8 min. Each column was then placed in a clean 1.5 mL tube and DNA was eluted by adding 700 µL of elution buffer (10 mM Tris-HCl pH 8.0) to the membrane and incubating at 22 °C for 2 min, followed by centrifugation at 12,000 g for 2 min. DNA concentration was determined with a Qubit 2.0 fluorometer with dsDNA HS chemistry (Thermo Scientific) and diluted to 0.01 ng/µL for PCR.

## Supplementary Figures

**Figure S1.**
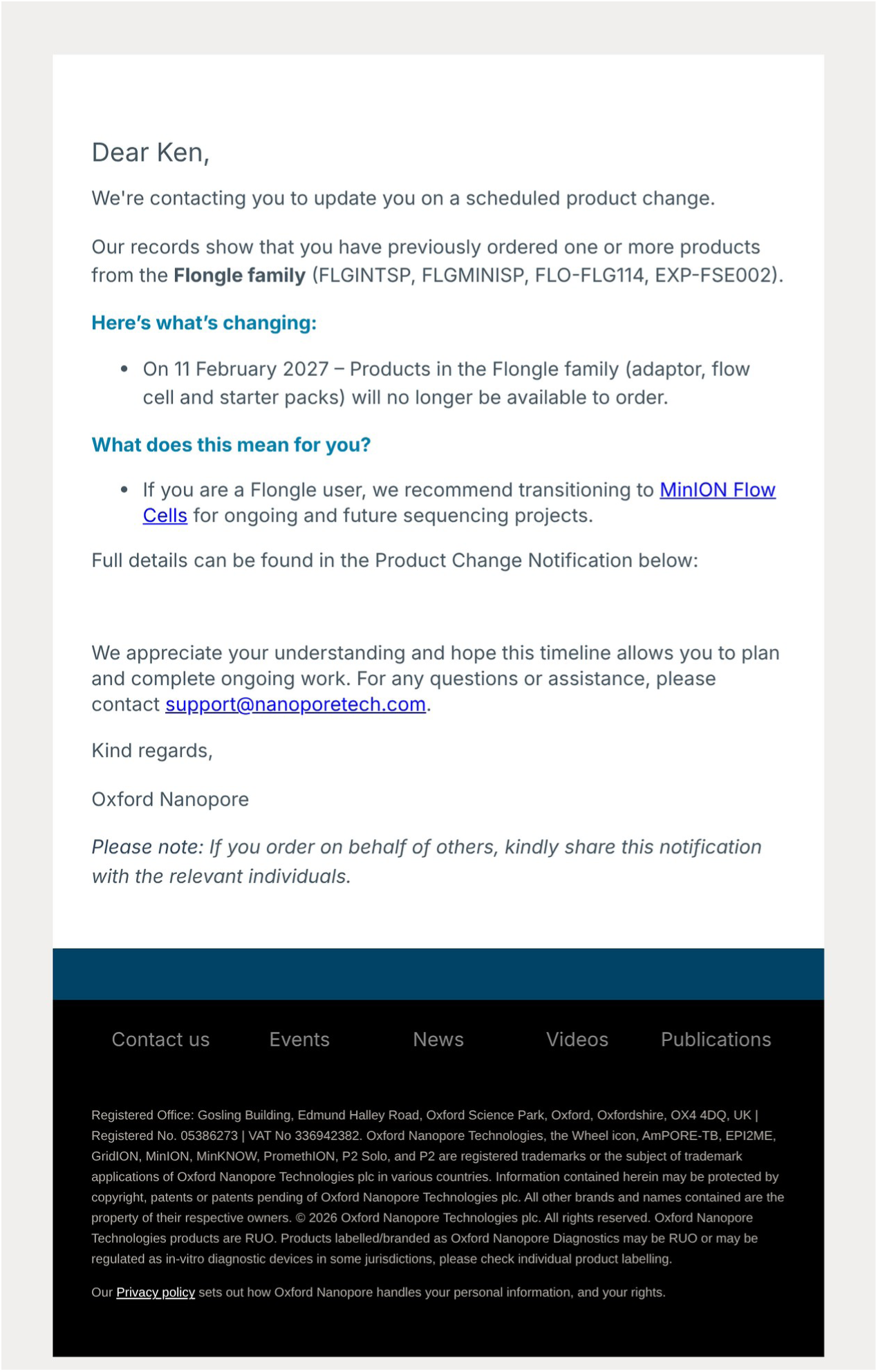
Product change notification from Oxford Nanopore Technologies (email, 2026): products in the Flongle family (adaptor, flow cell and starter packs), used for the ONT runs in Experiment 2, can no longer be ordered after 11 February 2027; ONT recommends MinION flow cells instead.

**Figure S2.**
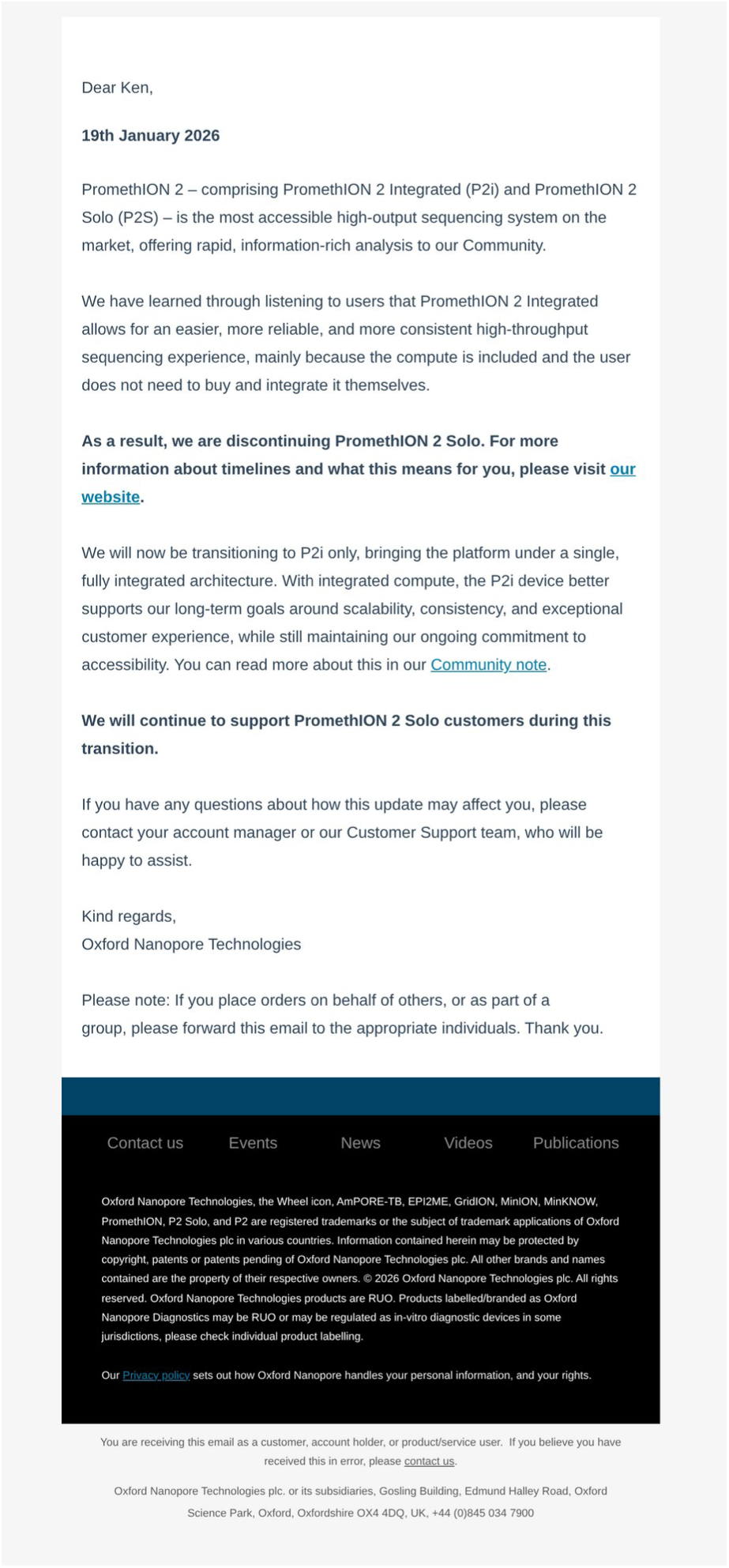
Notification from Oxford Nanopore Technologies (email, 19 January 2026) that the PromethION 2 Solo (P2S) is discontinued, with the platform moving to the PromethION 2 Integrated (P2i) only.

**Figure S3.**
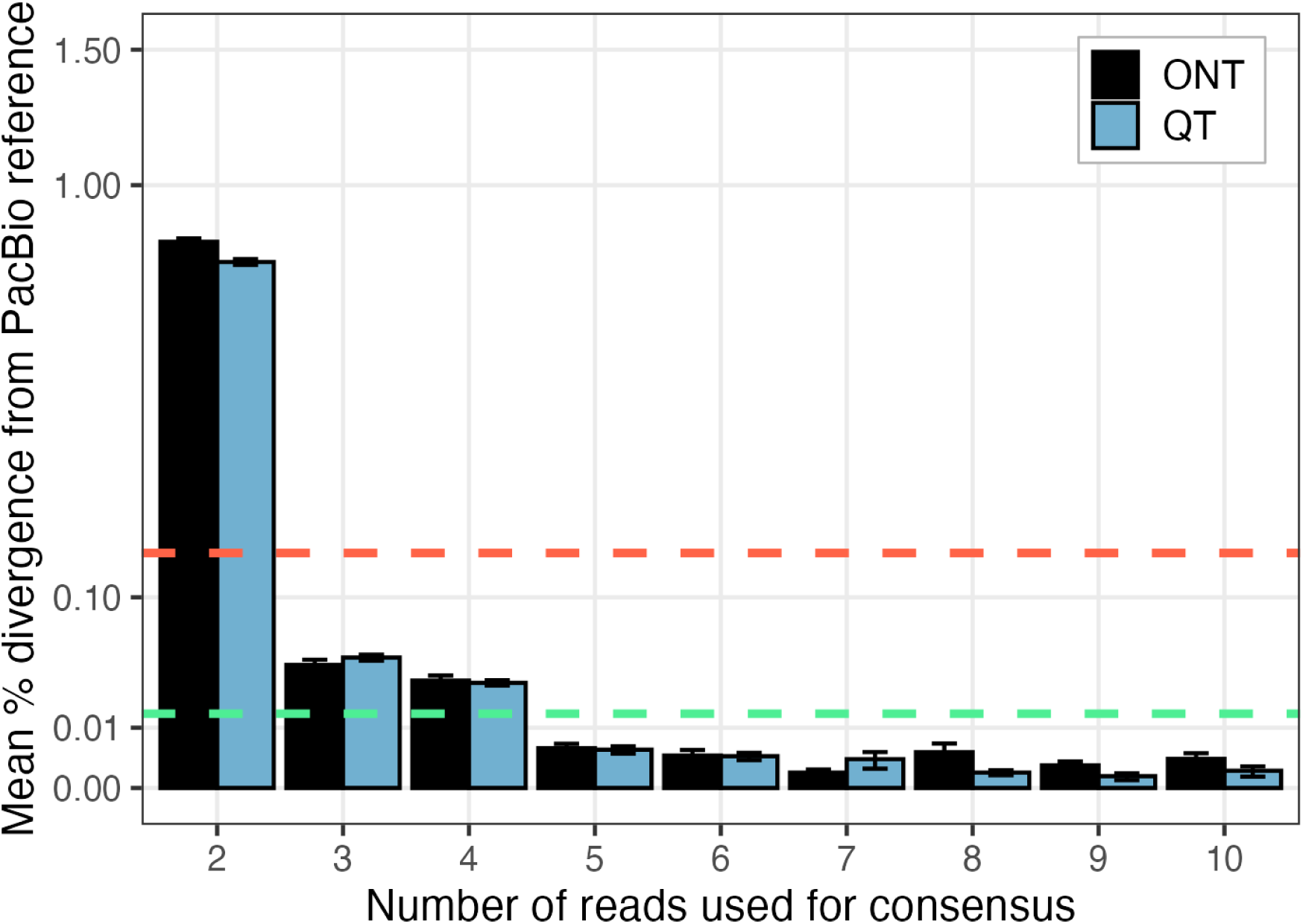
Evaluation of read accuracy for ONT (Flongle; Black) and Qitan (QCell; Blue) in Experiment 2, as in Figure 3B. Analysis included all 1,398 specimens where ONT, QT and the specimen’s gold-standard BOLD (PacBio SMRT) sequence were identical (658 bp, no ambiguous bases), with at least 10 reads supporting the barcode on each platform. The mean % sequence distance from the PacBio reference is shown for varying numbers of underlying sequences. The y- axis is on a square-root scale. The red dashed line reflects one error in a 658 bp sequence; the green dashed line reflects one error in ten 658 bp sequences. Error bars represent ±1 SE.

**Figure S4.**
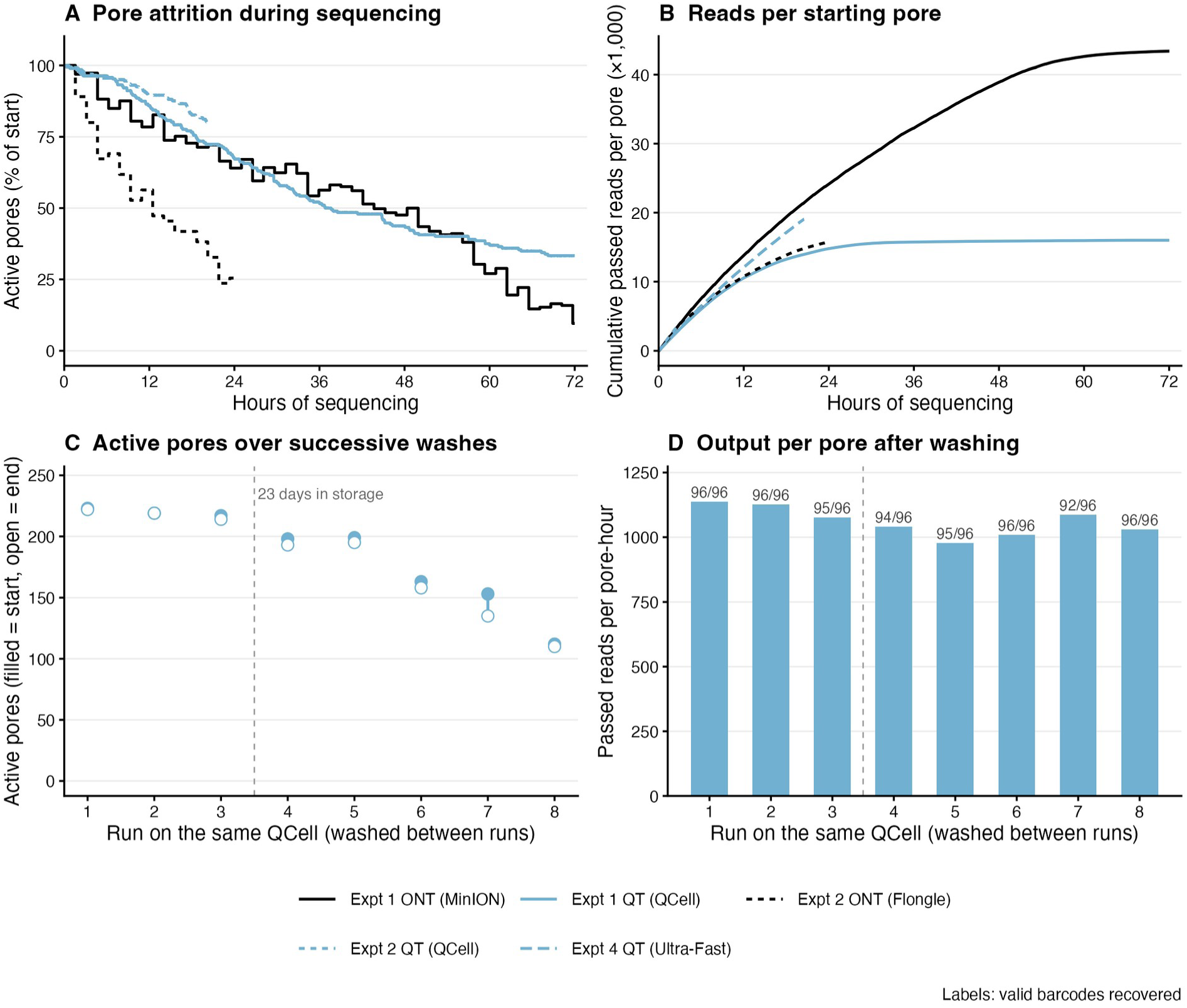
Pore attrition and output per pore. (A) Active pores during each main run as a percentage of those active at the start (ONT: single-pore channels at each mux scan, every ∼1.5 h; QT: available channels minus those the QNome reports as failed, every 30 s). (B) Cumulative passed reads per pore active at the start. (C) Active pores at the start (filled) and end (open) of eight successive runs on one QCell (Experiment 3), washed between runs; runs 1–3 and 4–8 were separated by 23 days in storage. (D) Passed reads per pore-hour in the same runs, with the valid barcodes recovered from each run’s 96-well plate.

**Figure S5.**
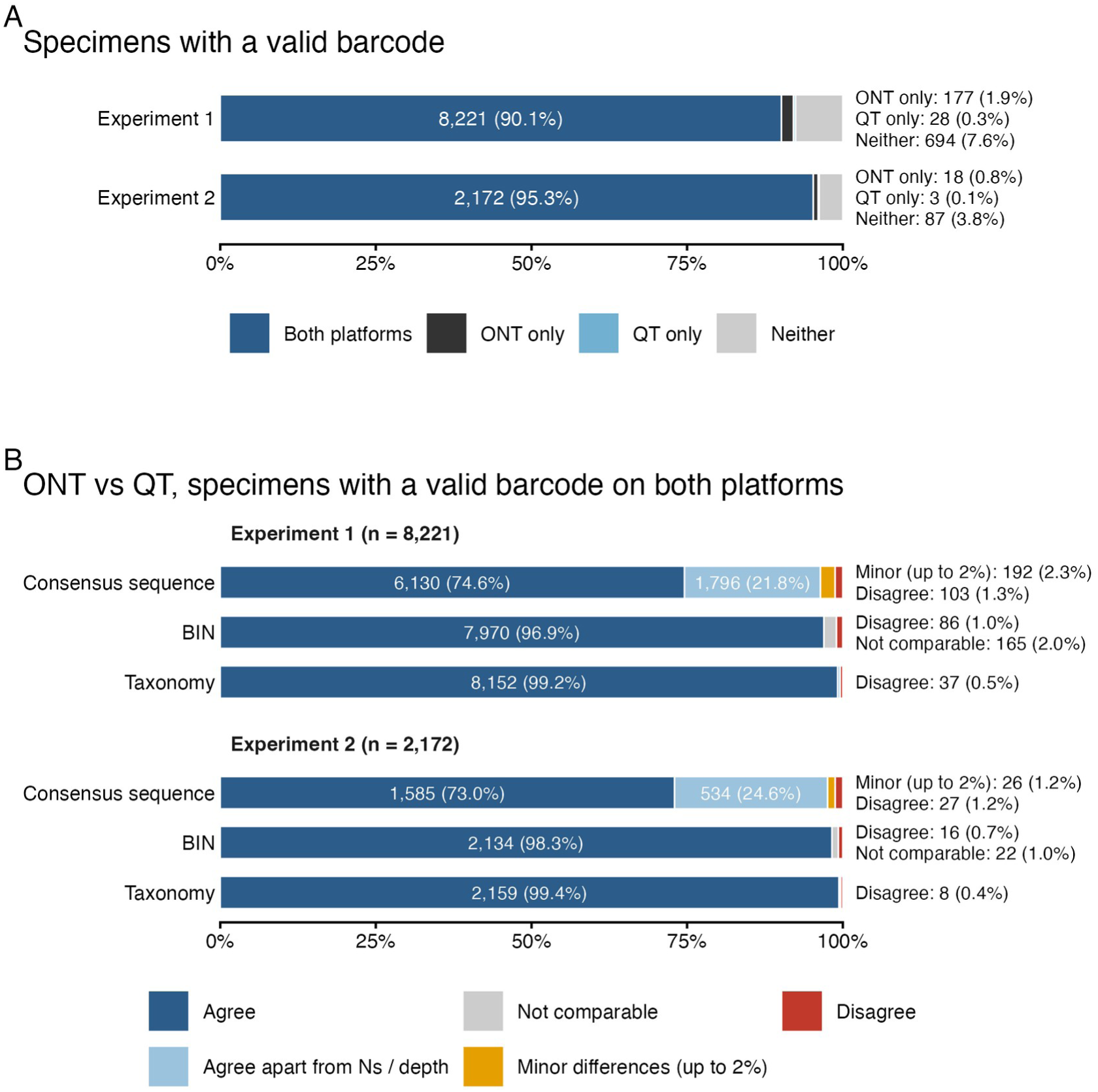
Visualization of the data in Tables S6 and S7. (A) Valid barcode recovery per specimen in Experiments 1 and 2: recovered by both platforms, ONT only, QT only, or neither (Table S6). (B) For specimens with a valid barcode on both platforms, agreement of the consensus sequence, BIN and taxonomic assignment between ONT and QT (Table S7). “Agree apart from Ns / depth” covers sequences that differ only at Ns and taxonomy where one platform is more resolved; For BINs, “Disagree” includes specimens assigned a BIN on one platform only, and “Not comparable” covers specimens with no BIN on either platform; for consensus sequences, “Disagree” means more than 2% of positions differ. Segments under 8% are labelled to the right of the bar.

**Figure S6.**
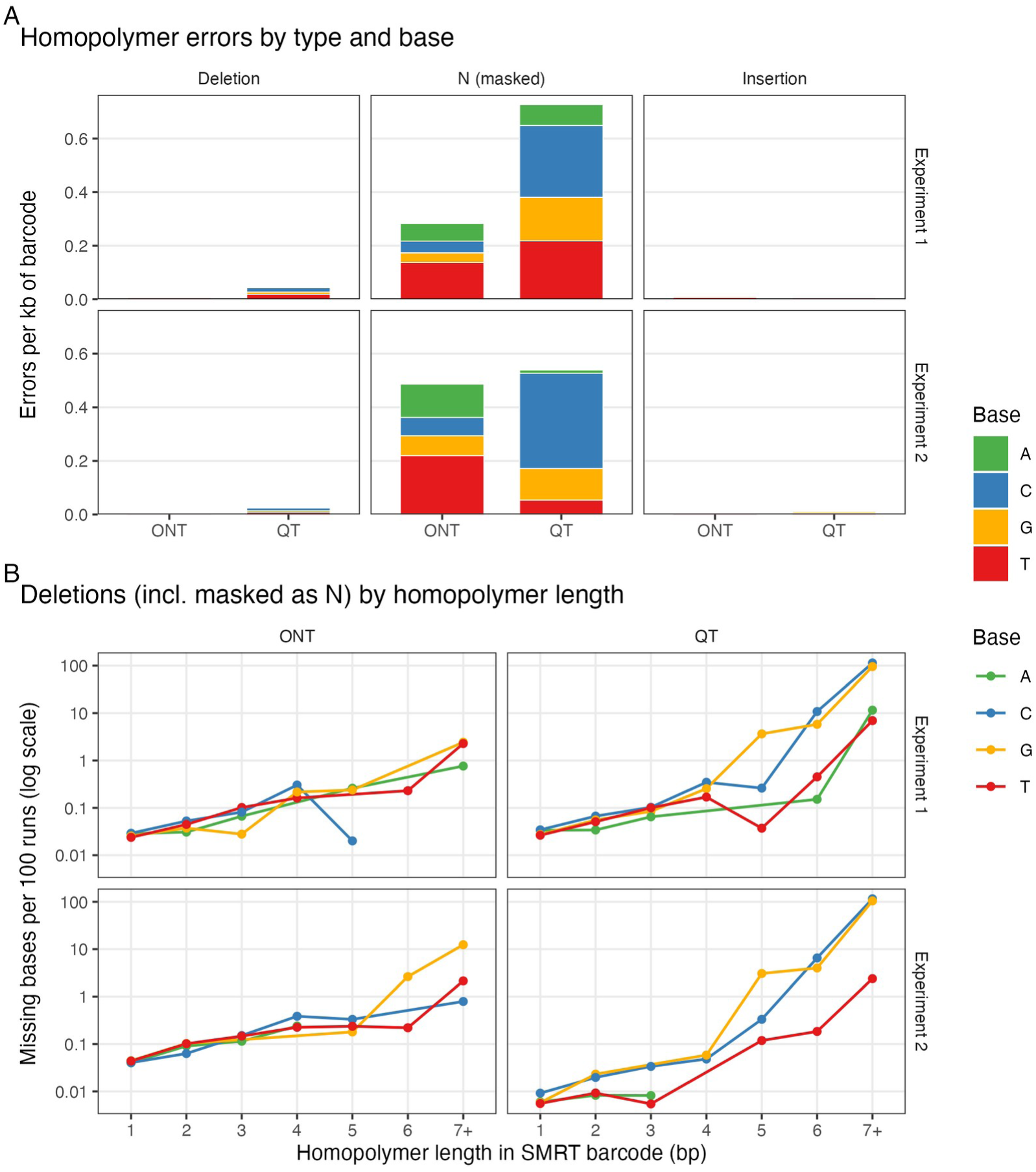
Homopolymer errors in ONT and QT barcodes, using each specimen’s PacBio SMRT barcode in BOLD (Experiment 1: DS-QT96; Experiment 2: DS-QT24) as the reference. For each specimen and platform, BIP’s target OTU was used, or, if it did not qualify, the closest NTS OTU; a sequence qualified if its assigned order matched the BOLD record and it differed from the SMRT barcode at no more than 5% of positions (Experiment 1: ONT n = 8,015, QT n = 8,002; Experiment 2: ONT n = 2,088, QT n = 2,086). Sequences were aligned to the SMRT barcode without end-gap penalties and every indel was treated as a homopolymer error. (A) Errors per kb of barcode by type and by the base involved: deletions (a base missing), Ns where SMRT has a base (BIP’s homopolymer correction inserts an N for a missing base, so these are masked deletions), and insertions (an extra base). BIP also removes extra bases when correcting, so insertions remaining in final barcodes underestimate those produced by sequencing. (B) Missing bases (deletions and masked deletions) per 100 homopolymer runs of each base and length in the SMRT barcodes; values above 100 mean runs lost more than one base. Points with no missing bases are not shown on the log scale.

**Figure S7.**
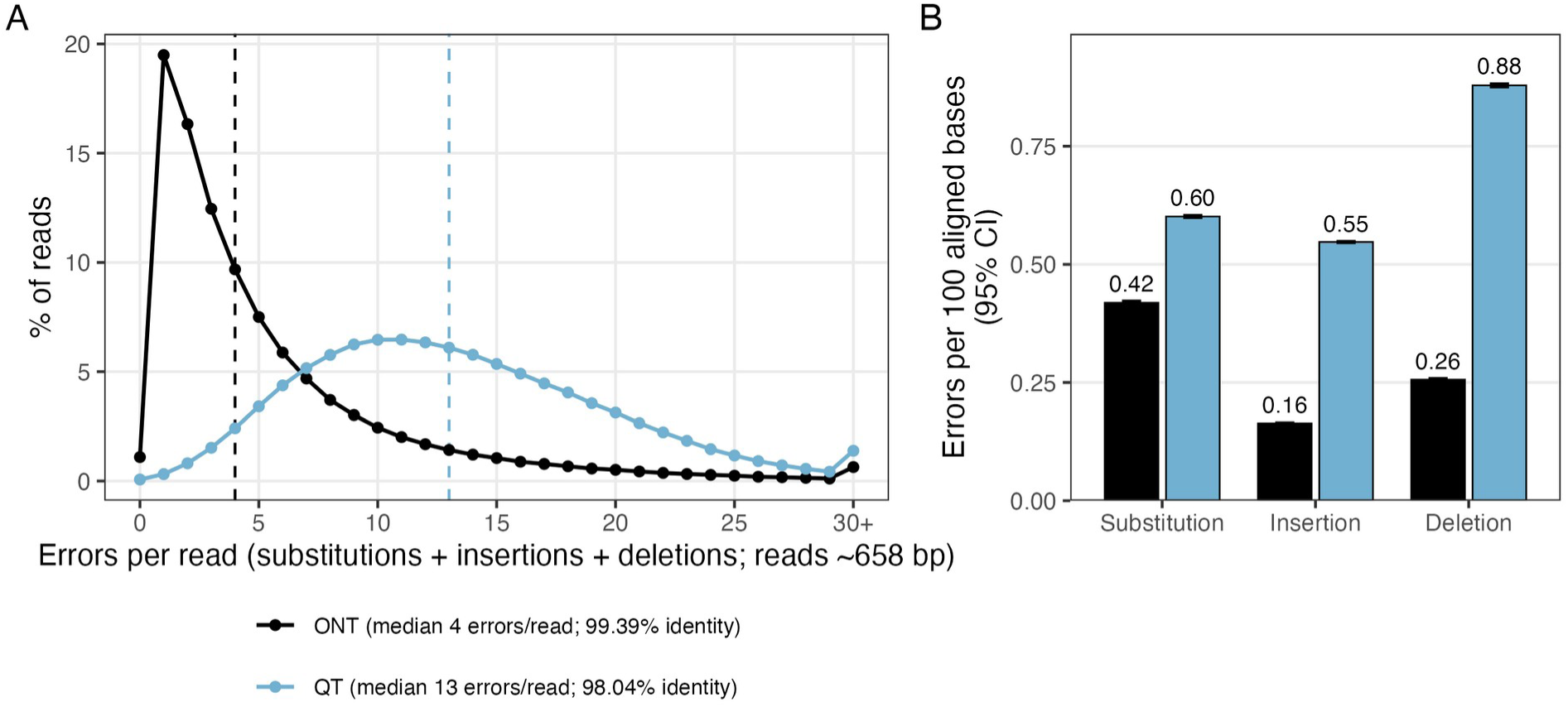
Per-read accuracy of ONT (MinION; black) and QT (QCell; blue) in Experiment 1. Every read in each platform’s barcode OTU for the 5,399 specimens of Figure 3B (ONT, QT and the gold-standard BOLD PacBio SMRT sequence identical, 658 bp) was aligned with minimap2 to that specimen’s sequence. (A) Errors per read (substitutions + insertions + deletions); dashed lines mark the medians. (B) Substitutions, insertions and deletions per 100 aligned bases, with 95% confidence intervals from a bootstrap over specimens (B = 2,000). Values for every metric are in Table S11.

**Figure S8.**
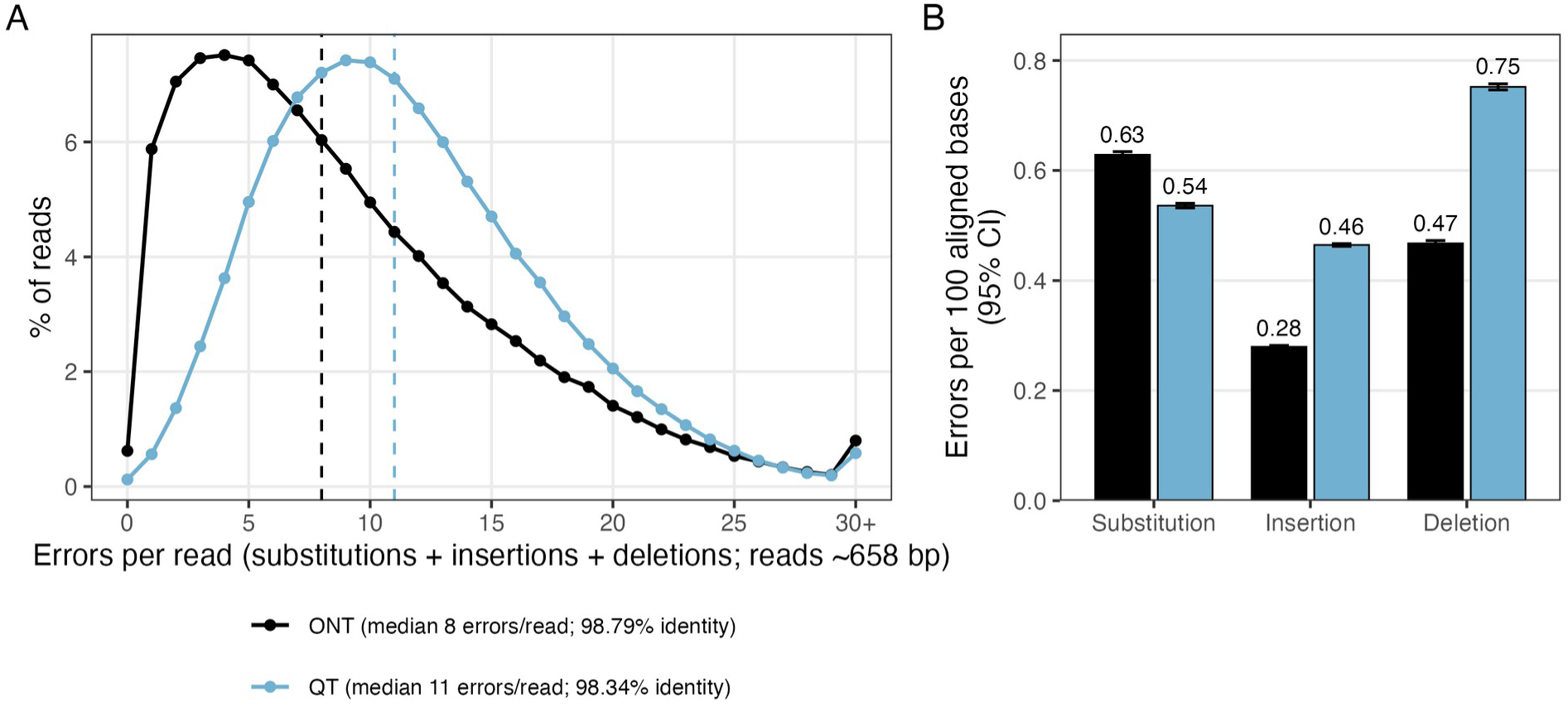
Per-read accuracy of ONT (Flongle; black) and QT (QCell; blue) in Experiment 2. Every read in each platform’s barcode OTU for the 1,398 specimens of Figure S3 (ONT, QT and the gold- standard BOLD PacBio SMRT sequence identical, 658 bp) was aligned with minimap2 to that specimen’s sequence. (A) Errors per read (substitutions + insertions + deletions); dashed lines mark the medians. (B) Substitutions, insertions and deletions per 100 aligned bases, with 95% confidence intervals from a bootstrap over specimens (B = 2,000). Values for every metric are in Table S12.

**Figure S9.**
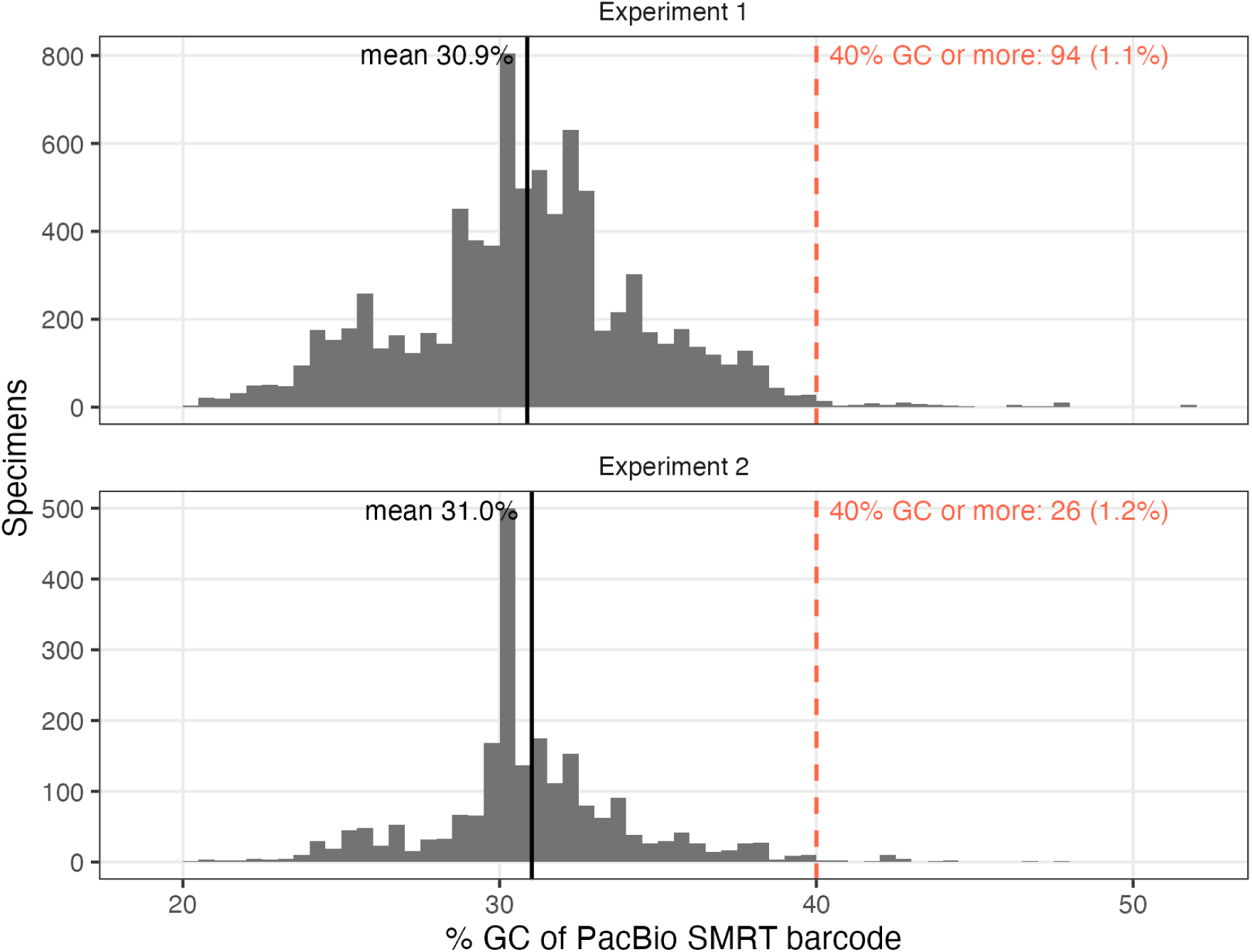
Distribution of barcode GC content per specimen in Experiments 1 and 2, from each specimen’s PacBio SMRT barcode in BOLD (Table S13). Black lines mark the mean; the red dashed line marks 40% GC. Across the 9,322 unique specimens, mean GC was 30.9% (median 30.9%) and 109 (1.2%) had barcodes of 40% GC or more.

## Supplementary Tables

**Table S1.** Experiment 2 (Flongle vs QCell) with every Q ≥ 10 read. The first two columns are Table 2, where the QCell reads were downsampled to the number of Flongle reads; the third column gives the QCell result using all its reads. Reads analysed is the number of reads (mean Q ≥ 10) given to BIP. Percentages of ambiguous-base OTUs are of valid barcodes. All runs used BIP v1.1.0 with the settings of the main analyses; the full-depth runs differ only in that no reads were discarded to equalise depth.

|  | *ONT (Flongle), as Table 2* | *QT (QCell), as Table 2* | *QT (QCell), all reads* |
|---|---|---|---|
| *Reads analysed (mean Q ≥ 10)* | 859,872 | 859,872 | 891,015 |
| *Reads in OTUs (% retained)* | 519,977 (60.5%) | 486,378 (56.6%) | 503,582 (56.5%) |
| *# Valid Barcodes (% success)* | 2,190 (96.1%) | 2,175 (95.4%) | 2,176 (95.4%) |
| *Valid barcodes with ambiguous bases* | 88 (4.0%) | 596 (27.4%) | 589 (27.1%) |
| *Mean read depth of barcodes (±SD)* | 225.9 (±84.4) | 211.1 (±76.4) | 218.5 (±79.1) |

**Table S2.** Experiment 4 (QCell standard vs ultra-fast library prep) with every Q ≥ 10 read. The first two columns are Table 4, where both libraries were given the same number of reads; the last two columns use all reads of each run. The ultra-fast library retained few reads through demultiplexing (25% of reads ended in OTUs vs 56% for the standard library), so equal input gives it a lower barcode depth. Reads analysed is the number of reads (mean Q ≥ 10) given to BIP. Percentages of ambiguous-base OTUs are of valid barcodes. All runs used BIP v1.1.0 with the settings of the main analyses; the full-depth runs differ only in that no reads were discarded to equalise depth.

|  | *Standard, as Table 4* | *Ultra-Fast, as Table 4* | *Standard, all reads* | *Ultra-Fast, all reads* |
|---|---|---|---|---|
| *Reads analysed (mean Q ≥ 10)* | 859,872 | 859,872 | 891,015 | 3,848,241 |
| *Reads in OTUs (% retained)* | 486,378 (56.6%) | 217,133 (25.3%) | 503,582 (56.5%) | 958,322 (24.9%) |
| *# Valid Barcodes (% success)* | 2,175 (95.4%) | 2,126 (93.2%) | 2,176 (95.4%) | 2,173 (95.3%) |
| *Valid barcodes with ambiguous bases* | 596 (27.4%) | 655 (30.8%) | 589 (27.1%) | 582 (26.8%) |
| *Mean read depth of barcodes (±SD)* | 211.1 (±76.4) | 90.7 (±38.2) | 218.5 (±79.1) | 374.6 (±175.9) |

**Table S3.** Experiment 1 (MinION vs QCell) with every Q ≥ 10 read. The first and last columns are Table 1, where MinION reads were downsampled to the number of QCell reads; the middle column gives the MinION result using all its reads. Reads analysed is the number of reads (mean Q ≥ 10) given to BIP. Percentages of ambiguous-base OTUs are of valid barcodes. All runs used BIP v1.1.0 with the settings of the main analyses; the full-depth runs differ only in that no reads were discarded to equalise depth.

|  | *ONT (MinION), as Table 1* | *ONT (MinION), all reads* | *QT (QCell), as Table 1* |
|---|---|---|---|
| *Reads analysed (mean Q ≥ 10)* | 3,070,864 | 21,111,283 | 3,070,864 |
| *Reads in OTUs (% retained)* | 2,035,597 (66.3%) | 14,060,174 (66.6%) | 1,512,516 (49.3%) |
| *# Valid Barcodes (% success)* | 8,398 (92.1%) | 8,528 (93.5%) | 8,249 (90.4%) |
| *Valid barcodes with ambiguous bases* | 335 (4.0%) | 265 (3.1%) | 2,055 (24.9%) |
| *Mean read depth of barcodes (±SD)* | 222.3 (±97.2) | 1,503.6 (±695.2) | 163.1 (±69.0) |

**Table S4.** Sequencing output per active pore for every run, from the instrument logs (QT: QNome sequence and flow-cell-check reports; ONT: MinKNOW throughput, pore-scan and sequencing- summary files). Active pores: QT, channels available at the flow-cell check and at the end of the run (QNome report); ONT, channels with a single pore in at least one well at the first and last mux scan (a MinION channel sequences one of its four wells at a time, so channels are the unit equivalent to a QCell channel; 1,168 of 2,048 MinION wells had a single pore at the start). Pore- hours: the area under the active-pore curve (Figure S4A). Passed reads and bases use each instrument’s own filter (MinKNOW pass; QNome Q10, or Q7 for Expt 2; Expt 4 was re-basecalled offline at Q10); Q≥10 reads use the filter applied before BIP (mean error probability). Rows from reads per pore-hour down measure platform capacity; the first 2.8 h compares every run over the length of the shortest one. QT Expt 2 uses all 891,015 Q≥10 reads (2,176 valid barcodes; 2,175 at the equal depth of Table 2). Wash runs (Expt 3) were stopped at about 100,000 reads and each sequenced one 96-well plate; values are the mean (range) of the eight runs and barcodes are those on the run’s own plate.

| *Metric* | *Expt 1 ONT (MinION)* | *Expt 1 QT (QCell)* | *Expt 2 ONT (Flongle)* | *Expt 2 QT (QCell)* | *Expt 4 QT (QCell, Ultra-Fast)* | *Expt 3 QT wash runs 1–8, mean (range)* |
|---|---|---|---|---|---|---|
| *Flow cell* | MinION | QCell | Flongle | QCell | QCell | QCell |
| *Run time (h)* | 72.0 | 72.0 | 24.0 | 2.9 | 20.5 | 0.6 (0.4–0.9) |
| *Active pores, start* | 492 | 192 | 55 | 261 | 202 | 186 (112–223) |
| *Active pores, end (% lost; wash runs: % lost)* | 47 (90%) | 64 (67%) | 14 (75%) | 254 (3%) | 162 (20%) | 3 (0–12) |
| *Pore-hours* | 19,960 | 8,129 | 735 | 740 | 3,797 | 101 (91–113) |
| *Reads passing instrument filter (M)* | 21.34 | 3.07 | 0.87 | 1.00 | 3.85 | 0.11 (0.10–0.11) |
| *Bases passing instrument filter (Gb)* | 16.63 | 2.36 | 0.67 | 0.78 | 2.91 | 0.08 (0.08–0.08) |
| *Reads per active pore* | 43,384 | 16,001 | 15,750 | 3,845 | 19,059 | 601 (464–904) |
| *Hours to 90% of passed reads* | 48.3 | 22.3 | 18.8 | 2.6 | 18.1 | 0.6 (0.4–0.9) |
| *Reads per pore-hour, whole run* | 1,069 | 378 | 1,178 | 1,356 | 1,014 | 1,061 (978–1,137) |
| *Reads per pore-hour, first 2.8 h* | 1,325 | 1,065 | 1,275 | 1,357 | 1,102 | – |
| *kb per pore-hour* | 832.9 | 290.8 | 913.1 | 1,050.7 | 765.9 | 808.4 (740.9–874.5) |
| *Q≥10 reads per active pore* | 42,909 | 15,994 | 15,634 | 3,414 | 19,051 | 598 (462–899) |
| *Reads analysed by BIP (% of Q≥10)* | 100.0 | 100.0 | 100.0 | 100.0 | 100.0 | 94.4 (90.4–99.3) |
| *Reads in OTUs per active pore* | 28,578 | 7,878 | 9,454 | 1,929 | 4,744 | 293 (244–457) |
| *Valid barcodes (of specimens)* | 8,528 / 9,120 | 8,249 / 9,120 | 2,190 / 2,280 | 2,176 / 2,280 | 2,173 / 2,280 | 95 (92–96) |
| *Valid barcodes per active pore* | 17.33 | 42.96 | 39.82 | 8.34 | 10.76 | 0.54 (0.43–0.86) |
| *Valid barcodes per sequencing hour* | 118 | 115 | 91 | 761 | 106 | 174 (108–238) |

**Table S5.**
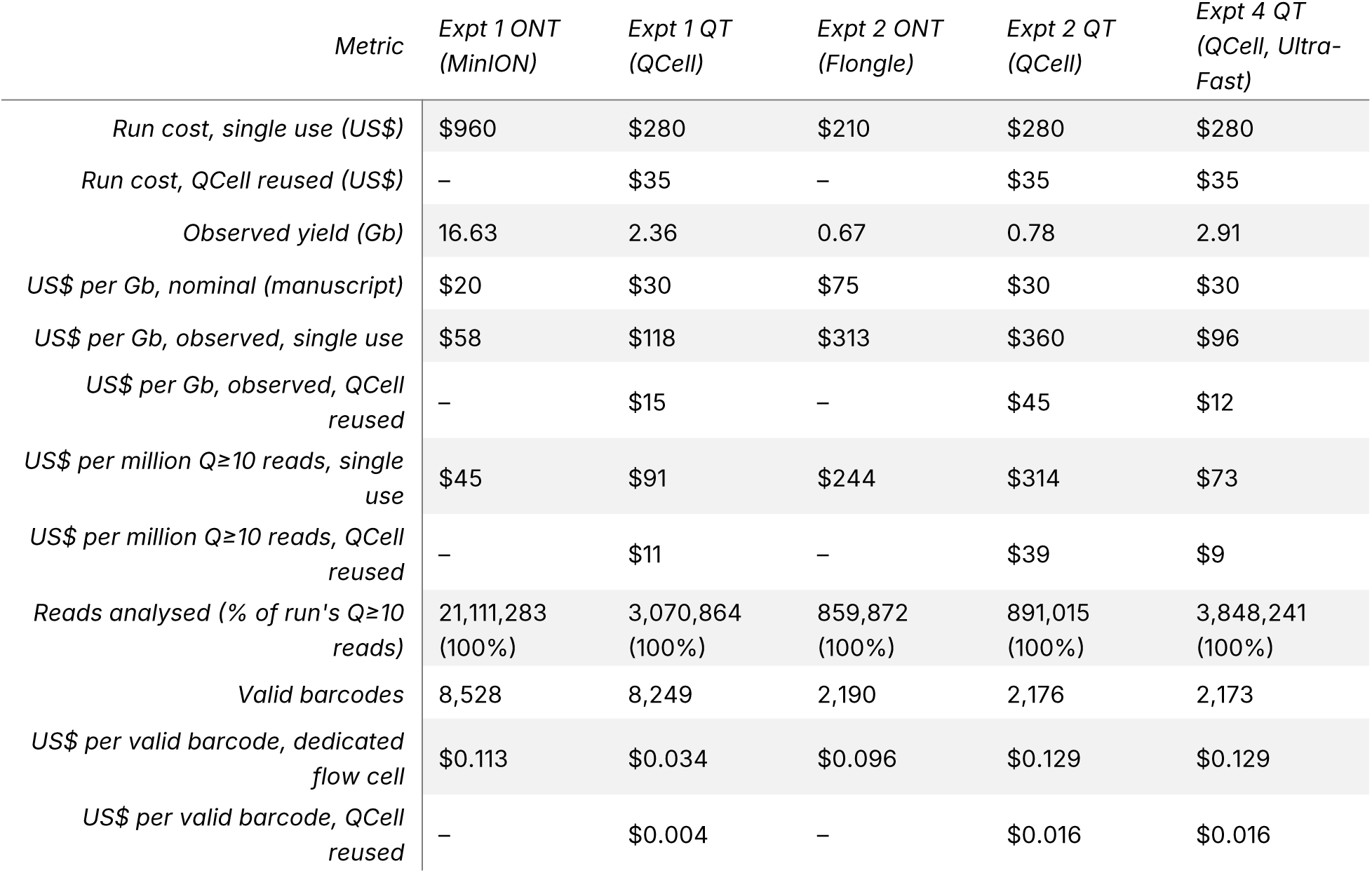
Consumable cost per run and per unit of output. Prices are those given in the Methods: Flongle flow cell US$210 (including reagents, single use), MinION flow cell US$960, QCell US$280 (including reagents). Instrument purchase prices (QNome and MinION) are excluded; the QNome price cannot be disclosed at QT’s request. Other costs not included: ONT library kits, QCell washes, ONT flow-cell washes, QCell shipping and brokerage, ONT shipping. Leaving out the instrument prices and ONT library kits understates costs; leaving out wash and shipping costs understates QT costs, so “QCell reused” is a lower bound. “QCell reused” divides the QCell price over the 8 runs one QCell delivered in Experiment 3 (≥ 92 of 96 barcodes in every run); these were eight runs of about 100,000 reads each, so the “QCell reused” figures assume runs of that size. MinION flow cells can also be washed but reuse was not tested here, so MinION is costed as single use. Observed yields are passed bases (Table S4) and replace the nominal yields behind the per-Gb costs in the Methods (shown for comparison). Per-barcode costs use the barcodes BIP recovered from the reads it analysed, all of a run’s Q≥10 reads: “dedicated flow cell” charges the whole run to those barcodes. The amplicon PCR upstream of library preparation is identical for both platforms and is excluded.

| *Metric* | *Expt 1 ONT (MinION)* | *Expt 1 QT (QCell)* | *Expt 2 ONT (Flongle)* | *Expt 2 QT (QCell)* | *Expt 4 QT (QCell, Ultra-Fast)* |
|---|---|---|---|---|---|
| *Run cost, single use (US\$)* | \$960 | \$280 | \$210 | \$280 | \$280 |
| *Run cost, QCell reused (US\$)* | – | \$35 | – | \$35 | \$35 |
| *Observed yield (Gb)* | 16.63 | 2.36 | 0.67 | 0.78 | 2.91 |
| *US\$ per Gb, nominal (manuscript)* | \$20 | \$30 | \$75 | \$30 | \$30 |
| *US\$ per Gb, observed, single use* | \$58 | \$118 | \$313 | \$360 | \$96 |
| *US\$ per Gb, observed, QCell reused* | – | \$15 | – | \$45 | \$12 |
| *US\$ per million Q≥10 reads, single use* | \$45 | \$91 | \$244 | \$314 | \$73 |
| *US\$ per million Q≥10 reads, QCell reused* | – | \$11 | – | \$39 | \$9 |
| *Reads analysed (% of run's Q≥10 reads)* | 21,111,283 (100%) | 3,070,864 (100%) | 859,872 (100%) | 891,015 (100%) | 3,848,241 (100%) |
| *Valid barcodes* | 8,528 | 8,249 | 2,190 | 2,176 | 2,173 |
| *US\$ per valid barcode, dedicated flow cell* | \$0.113 | \$0.034 | \$0.096 | \$0.129 | \$0.129 |
| *US\$ per valid barcode, QCell reused* | – | \$0.004 | – | \$0.016 | \$0.016 |

**Table S6.**
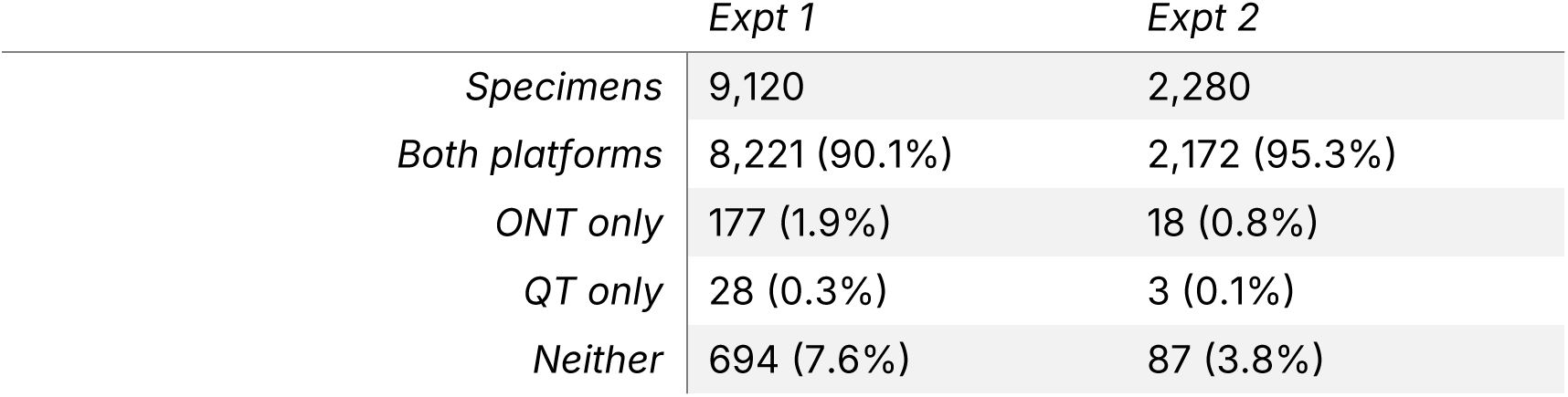
Valid barcode recovery per specimen by ONT and QT in Experiments 1 and 2. Each specimen (negative controls excluded) is counted once, by whether a valid barcode was recovered on both platforms, on one only, or on neither. Each experiment is counted separately: 1,330 specimens were sequenced in both experiments (matched by BOLD sample ID).

|  | *Expt 1* | *Expt 2* |
|---|---|---|
| *Specimens* | 9,120 | 2,280 |
| *Both platforms* | 8,221 (90.1%) | 2,172 (95.3%) |
| *ONT only* | 177 (1.9%) | 18 (0.8%) |
| *QT only* | 28 (0.3%) | 3 (0.1%) |
| *Neither* | 694 (7.6%) | 87 (3.8%) |

**Table S7.** Agreement between ONT and QT for specimens with a valid barcode on both platforms. Consensus sequences were globally aligned; a position differing only because one platform has an N was not counted as a difference. More than 13 differing bases (> 2% of 658 bp) indicates that the platforms recovered different sequences, i.e. a different organism was called as the specimen’s barcode. Taxonomy compares BIP’s assignments at order, family, genus and species: a conflict is a rank where both platforms assigned a name and the names differ, reported at the highest such rank; “more resolved” means no conflict but one platform assigned a lower rank. BIP assigns a BIN when the barcode matches a reference sequence at ≥ 97% identity over ≥ 75% of the amplicon; specimens with no BIN on either platform have no such reference. A BIN on one platform only is counted as a BIN disagreement: in 39 of 40 such specimens the consensus sequences were > 2% apart. Of all BIN disagreements (different BINs or a BIN on one platform only), 101 of 102 had consensus sequences > 2% apart, as did 45 of 45 specimens with a taxonomic conflict. Percentages are of specimens with a valid barcode on both platforms.

|  | *Expt 1* | *Expt 2* |
|---|---|---|
| *Specimens with a valid barcode on both platforms* | 8,221 | 2,172 |
| *Consensus sequence: identical* | 6,130 (74.6%) | 1,585 (73.0%) |
| *Consensus sequence: differ only at Ns* | 1,796 (21.8%) | 534 (24.6%) |
| *Consensus sequence: 1–13 bp differ (≤ 2%)* | 192 (2.3%) | 26 (1.2%) |
| *Consensus sequence: > 13 bp differ (> 2%)* | 103 (1.3%) | 27 (1.2%) |
| *BIN: same* | 7,970 (96.9%) | 2,134 (98.3%) |
| *BIN: different BINs* | 53 (0.6%) | 9 (0.4%) |
| *BIN: assigned on one platform only* | 33 (0.4%) | 7 (0.3%) |
| *BIN: not assigned on either platform* | 165 (2.0%) | 22 (1.0%) |
| *Taxonomy: same* | 8,152 (99.2%) | 2,159 (99.4%) |
| *Taxonomy: same, one platform more resolved* | 32 (0.4%) | 5 (0.2%) |
| *Taxonomy: conflict at order* | 0 (0.0%) | 0 (0.0%) |
| *Taxonomy: conflict at family* | 34 (0.4%) | 8 (0.4%) |
| *Taxonomy: conflict at genus* | 3 (< 0.1%) | 0 (0.0%) |
| *Taxonomy: conflict at species* | 0 (0.0%) | 0 (0.0%) |

**Table S8.**
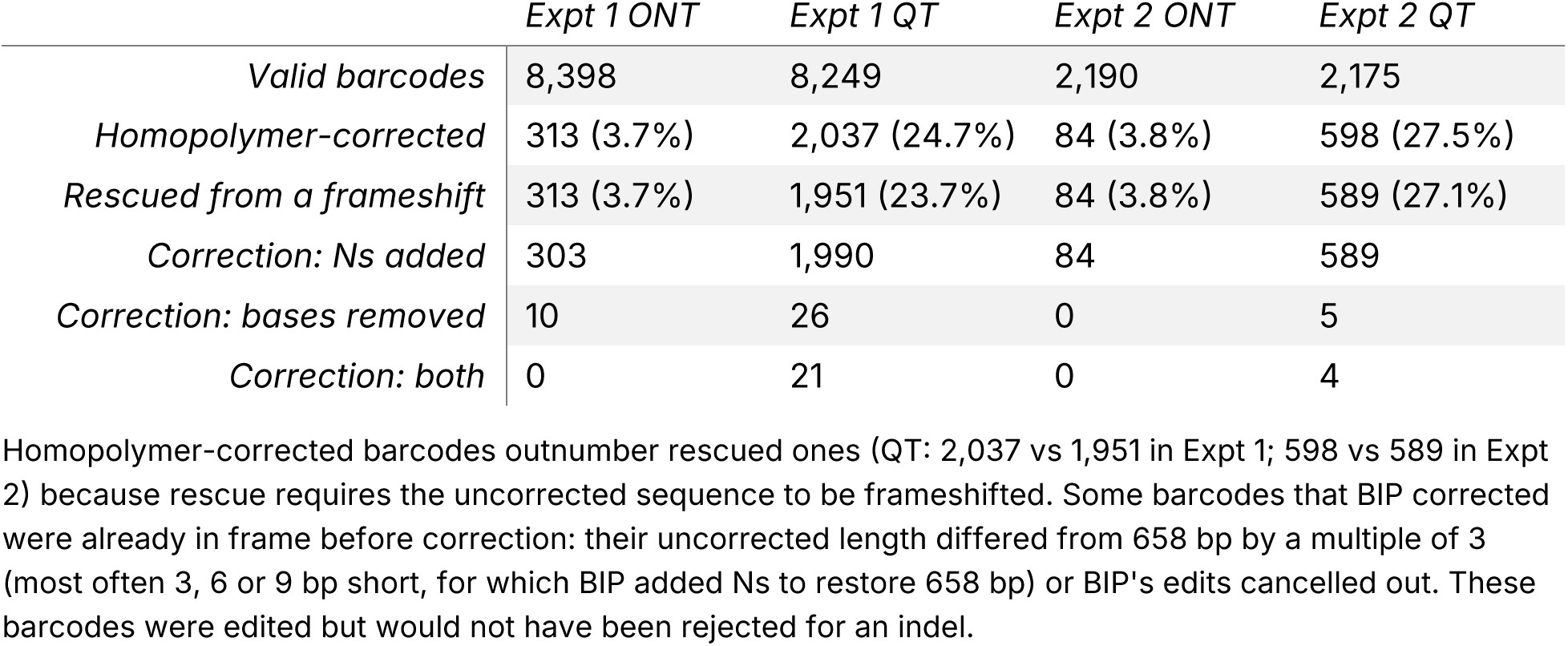
Valid barcodes that required BIP’s homopolymer error correction. A barcode was homopolymer-corrected when BIP added Ns where a base was missing in a homopolymer, or removed extra bases or Ns, to restore the reading frame against its closest reference sequence. It was rescued when the uncorrected sequence was frameshifted (length − 658 not a multiple of 3), so it would otherwise have been rejected for an indel. Percentages are of valid barcodes; the last three rows give the type of correction.

|  | *Expt 1 ONT* | *Expt 1 QT* | *Expt 2 ONT* | *Expt 2 QT* |
|---|---|---|---|---|
| *Valid barcodes* | 8,398 | 8,249 | 2,190 | 2,175 |
| *Homopolymer-corrected* | 313 (3.7%) | 2,037 (24.7%) | 84 (3.8%) | 598 (27.5%) |
| *Rescued from a frameshift* | 313 (3.7%) | 1,951 (23.7%) | 84 (3.8%) | 589 (27.1%) |
| *Correction: Ns added* | 303 | 1,990 | 84 | 589 |
| *Correction: bases removed* | 10 | 26 | 0 | 5 |
| *Correction: both* | 0 | 21 | 0 | 4 |

Homopolymer-corrected barcodes outnumber rescued ones (QT: 2,037 vs 1,951 in Expt 1; 598 vs 589 in Expt 2) because rescue requires the uncorrected sequence to be frameshifted. Some barcodes that BIP corrected were already in frame before correction: their uncorrected length differed from 658 bp by a multiple of 3 (most often 3, 6 or 9 bp short, for which BIP added Ns to restore 658 bp) or BIP's edits cancelled out. These barcodes were edited but would not have been rejected for an indel.

**Table S9.** Effect of QT homopolymer errors on barcode accuracy and identification, using ONT as the reference. Analysis is restricted to specimens whose ONT barcode was 658 bp with no ambiguous bases. Each QT OTU from these wells was aligned to the ONT barcode. “In or next to a homopolymer” means within, or 1 bp from, a run of ≥ 4 identical bases in the ONT barcode; the share of all barcode positions meeting this definition is shown for comparison. Percentages for QT barcode types are of QT valid barcodes; failure causes are of specimens without a valid QT barcode; other percentages are of specimens with a clean ONT barcode. BIN agreement counts barcodes assigned a BIN on both platforms. The last two rows give the QT barcode count if the closest QT OTU were corrected using the ONT sequence.

|  | *Expt 1* | *Expt 2* |
|---|---|---|
| *Specimens with a clean ONT barcode* | 7,471 | 2,019 |
| *QT valid barcode* | 7,351 (98.4%) | 2,006 (99.4%) |
| *QT barcode identical to ONT* | 5,677 (77.2%) | 1,477 (73.6%) |
| *QT barcode differs only at Ns* | 1,472 (20.0%) | 492 (24.5%) |
| *QT barcode differs at other positions* | 202 (2.7%) | 37 (1.8%) |
| *N bases in or next to a homopolymer* | 1,835 / 1,893 (96.9%) | 618 / 624 (99.0%) |
| *Barcode positions in or next to a homopolymer* | 16.5% | 16.3% |
| *Same BIN as ONT: QT barcodes with Ns* | 1,498 / 1,506 (99.5%) | 495 / 496 (99.8%) |
| *Same BIN as ONT: QT barcodes without Ns* | 5,694 / 5,731 (99.4%) | 1,489 / 1,496 (99.5%) |
| *No valid QT barcode* | 120 (1.6%) | 13 (0.6%) |
| *Failed on an uncorrected frameshift* | 85 (70.8%) | 8 (61.5%) |
| *QT barcodes if Ns replaced from ONT* | 7,354 (98.4%) | 2,006 (99.4%) |
| *QT barcodes if Ns and indels corrected from ONT* | 7,435 (99.5%) | 2,013 (99.7%) |

**Table S10.**
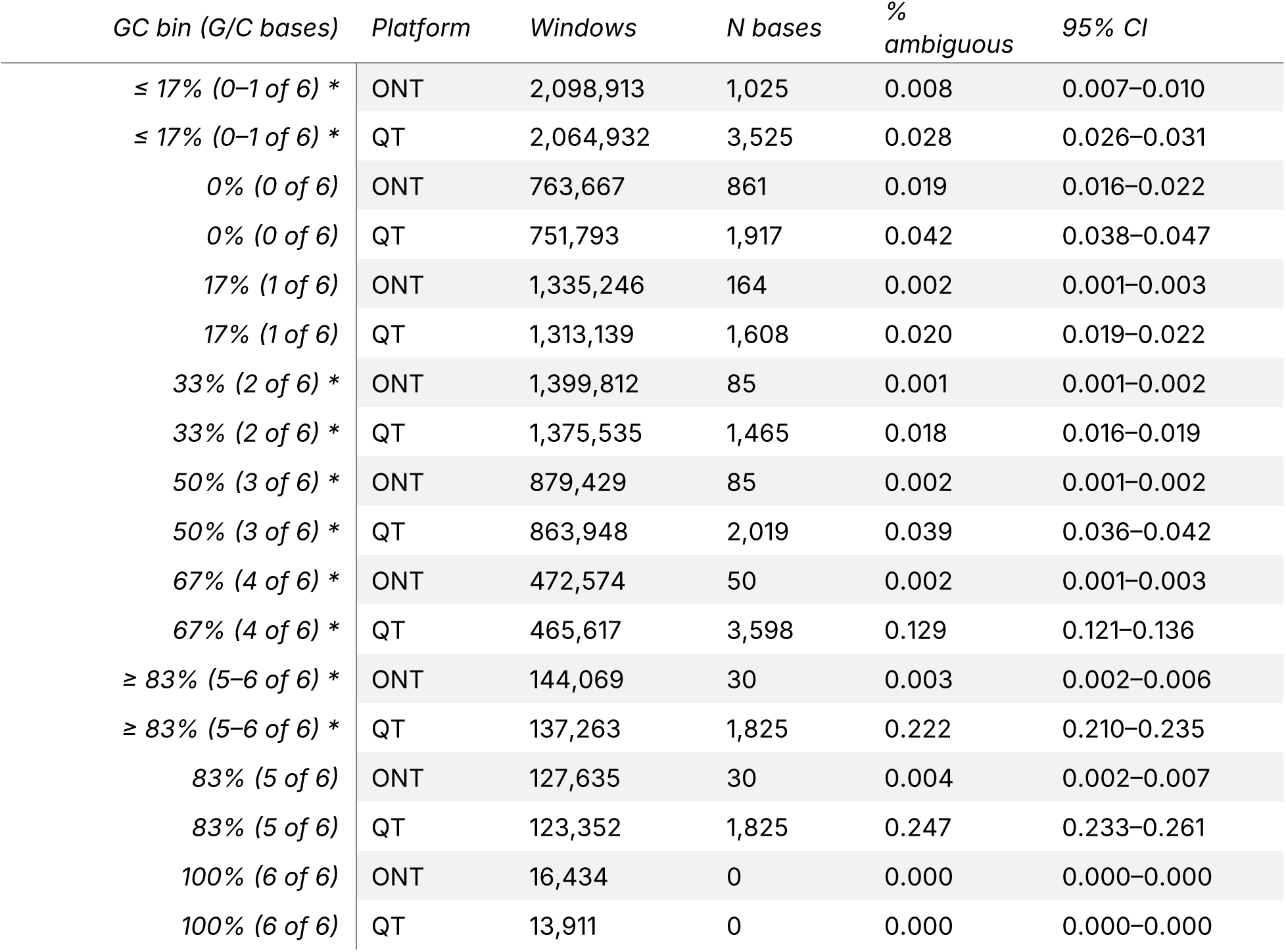
Ambiguous bases by GC content of 6 bp windows (Figure 3A): Experiment 1, valid 658 bp barcodes. Each overlapping 6 bp window (653 per barcode) was binned by its number of G or C bases (an N counts as neither). % ambiguous = N bases in the bin’s windows / all bases in those windows × 100; 95% CIs are percentiles of 2,000 bootstrap resamples of barcodes (within platform). Bins marked * are those plotted in Figure 3A, where both ends of the GC range are pooled; the single-count bins within them are listed beneath. Total windows: ONT 4,994,797, QT 4,907,295. A window containing an N has at most five G or C bases, so the 100% bin cannot contain an N; Ns at the end of G/C homopolymers are counted in the 83% bin.

**Table S11.**
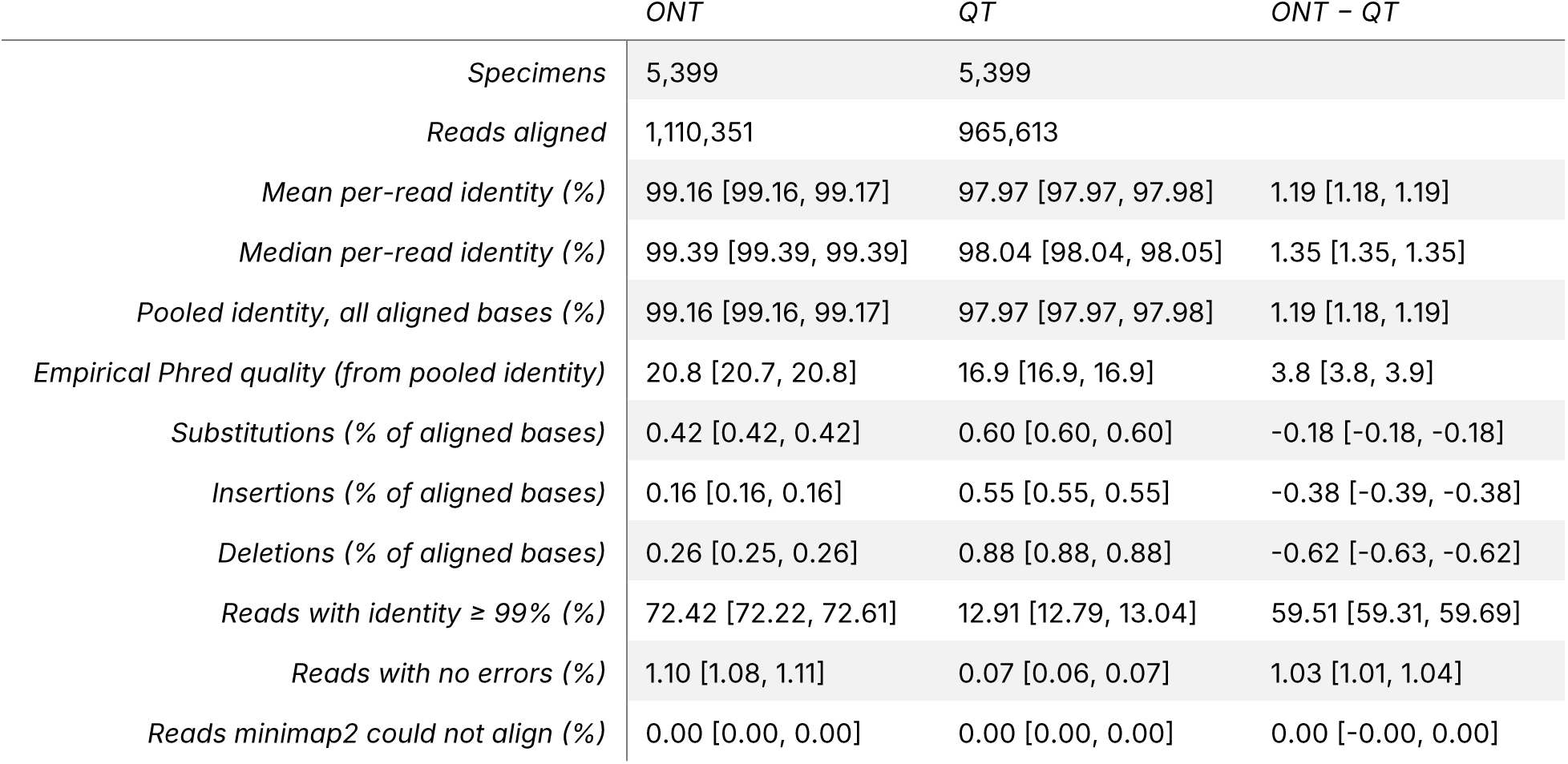
Per-read accuracy in Experiment 1 (5,399 specimens; reads as in Figure S7). Identity = matches / (matches + substitutions + insertions + deletions) in each read’s alignment to its specimen’s gold-standard reference; unaligned read ends are not counted. Values are pooled over reads, with 95% confidence intervals from a paired bootstrap over specimens (B = 2,000), which also gives the interval for the ONT − QT difference. Empirical Phred quality = −10·log10(1 − pooled identity). Only reads BIP clustered into the barcode OTU are included, so absolute accuracy is higher than for all reads in a run; both platforms were filtered identically (mean Q ≥ 10, same BIP settings). Some substitutions are PCR errors present on both platforms, so absolute accuracies are lower bounds on sequencing accuracy.

**Table S12.** Per-read accuracy in Experiment 2 (1,398 specimens; reads as in Figure S8). Identity = matches / (matches + substitutions + insertions + deletions) in each read’s alignment to its specimen’s gold-standard reference; unaligned read ends are not counted. Values are pooled over reads, with 95% confidence intervals from a paired bootstrap over specimens (B = 2,000), which also gives the interval for the ONT − QT difference. Empirical Phred quality = −10·log10(1 − pooled identity). Only reads BIP clustered into the barcode OTU are included, so absolute accuracy is higher than for all reads in a run; both platforms were filtered identically (mean Q ≥ 10, same BIP settings). Some substitutions are PCR errors present on both platforms, so absolute accuracies are lower bounds on sequencing accuracy.

|  | *ONT* | *QT* | *ONT − QT* |
|---|---|---|---|
| *Specimens* | 1,398 | 1,398 |  |
| *Reads aligned* | 328,317 | 318,798 |  |
| *Mean per-read identity (%)* | 98.63 [98.62, 98.64] | 98.25 [98.24, 98.26] | 0.38 [0.37, 0.39] |
| *Median per-read identity (%)* | 98.79 [98.79, 98.79] | 98.34 [98.34, 98.34] | 0.45 [0.45, 0.46] |
| *Pooled identity, all aligned bases (%)* | 98.62 [98.61, 98.63] | 98.25 [98.24, 98.26] | 0.38 [0.37, 0.39] |
| *Empirical Phred quality (from pooled identity)* | 18.6 [18.6, 18.6] | 17.6 [17.5, 17.6] | 1.1 [1.0, 1.1] |
| *Substitutions (% of aligned bases)* | 0.63 [0.62, 0.63] | 0.54 [0.53, 0.54] | 0.09 [0.09, 0.10] |
| *Insertions (% of aligned bases)* | 0.28 [0.28, 0.28] | 0.46 [0.46, 0.47] | -0.19 [-0.19, -0.18] |
| *Deletions (% of aligned bases)* | 0.47 [0.46, 0.47] | 0.75 [0.75, 0.76] | -0.28 [-0.29, -0.28] |
| *Reads with identity ≥ 99% (%)* | 42.94 [42.51, 43.34] | 19.09 [18.78, 19.40] | 23.86 [23.47, 24.22] |
| *Reads with no errors (%)* | 0.62 [0.60, 0.64] | 0.12 [0.11, 0.13] | 0.50 [0.48, 0.52] |
| *Reads minimap2 could not align (%)* | 0.01 [0.00, 0.01] | 0.00 [0.00, 0.00] | 0.00 [-0.00, 0.01] |

**Table S13.** GC content of the COI barcode per specimen in Experiments 1 and 2. GC = (G + C) / called bases × 100 (Ns and gaps ignored), from each specimen’s PacBio SMRT barcode in BOLD (Experiment 1: DS-QT96; Experiment 2: DS-QT24) and from the ONT and QT valid barcodes. The two experiments share 1,330 specimens, counted once in the combined row. The last two columns give the number (%) of specimens whose barcode GC is at or above 35% and 40%.

| *Experiment* | *Barcodes* | *Specimens* | *Mean % GC (SD)* | *Median (range)* | *≥ 35% GC* | *≥ 40% GC* |
|---|---|---|---|---|---|---|
| *Experiment 1* | PacBio SMRT (BOLD) | 8,388 | 30.9 (3.9) | 30.9 (19.4–51.8) | 1,093 (13.0%) | 94 (1.1%) |
| *Experiment 1* | ONT valid barcode | 8,398 | 30.9 (3.8) | 30.9 (19.4–51.8) | 1,114 (13.3%) | 74 (0.9%) |
| *Experiment 1* | QT valid barcode | 8,249 | 30.9 (3.8) | 30.9 (19.4–51.8) | 1,077 (13.1%) | 69 (0.8%) |
| *Experiment 2* | PacBio SMRT (BOLD) | 2,242 | 31.0 (3.3) | 30.4 (20.4–47.8) | 233 (10.4%) | 26 (1.2%) |
| *Experiment 2* | ONT valid barcode | 2,190 | 31.0 (3.3) | 30.5 (20.7–44.4) | 228 (10.4%) | 26 (1.2%) |
| *Experiment 2* | QT valid barcode | 2,175 | 31.0 (3.3) | 30.4 (20.7–44.4) | 232 (10.7%) | 26 (1.2%) |
| *Experiments 1 + 2 (unique specimens)* | PacBio SMRT (BOLD) | 9,322 | 30.9 (3.9) | 30.9 (19.4–51.8) | 1,205 (12.9%) | 109 (1.2%) |

## REFERENCES

1. Cameron, S. L. (2014). Insect mitochondrial genomics: implications for evolution and phylogeny. Annual Review of Entomology, 59(1), 95–117. 10.1146/annurev-ento-011613-162007

2. CBOL Plant Working Group. (2009). A DNA barcode for land plants. Proceedings of the National Academy of Sciences of the United States of America, 106(31), 12794–12797. 10.1073/pnas.0905845106

3. Dang, X., Ma, S., Jiang, J., Zhai, W., Cai, H., Lyu, H., Liao, Z., Yan, G., Wang, Z., Li, Y., Huang, J., Shen, J., Liu, D., Schwarz, S., Xia, Z., Wang, Y., Wang, L., & Chen, S. (2026). Antimicrobial resistance and transmission potential of pet- derived Providencia spp. in China. Journal of Global Antimicrobial Resistance, 48, 137–144. 10.1016/j.jgar.2026.03.004

4. Delahaye, C., & Nicolas, J. (2021). Sequencing DNA with nanopores: Troubles and biases. PloS One, 16(10), e0257521. 10.1371/journal.pone.0257521

5. Edgar, R. (2016). SINTAX: a simple non-Bayesian taxonomy classifier for 16S and ITS sequences. bioRxiv. 10.1101/074161

6. Edgar, R. C., Haas, B. J., Clemente, J. C., Quince, C., & Knight, R. (2011). UCHIME improves sensitivity and speed of chimera detection. Bioinformatics, 27(16), 2194–2200. 10.1093/bioinformatics/btr381

7. Elbrecht, V., Braukmann, T. W. A., Ivanova, N. V., Prosser, S. W. J., Hajibabaei, M., Wright, M., Zakharov, E. V., Hebert, P. D. N., & Steinke, D. (2019). Validation of COI metabarcoding primers for terrestrial arthropods. PeerJ, 7, e7745. 10.7717/peerj.7745

8. Floyd, R., Prosser, S., & Jafarpour, S. (2023). DNA barcoding on Oxford Nanopore: multiplexing up to 24 × 96-well plates. protocols.io. 10.17504/protocols.io.rm7vzx6n4gx1/v1

9. Frey, K., & Pucker, B. (2020). Animal, fungi, and plant genome sequences harbor different non-canonical splice sites. Cells (Basel, Switzerland), 9(2), 458. 10.3390/cells9020458

10. Gygax, D., Ramirez, S., Chibesa, M., Simpamba, T., Riffel, M., Riffel, T., Srivathsan, A., Nijland, R., & Urban, L. (2025). Evaluation of nanopore sequencing for increasing accessibility of eDNA studies in biodiverse countries. PloS One, 20(10), e0333994. 10.1371/journal.pone.0333994

11. Hebert, P. D. N., Braukmann, T. W. A., Prosser, S. W. J., Ratnasingham, S., deWaard, J. R., Ivanova, N. V., Janzen, D. H., Hallwachs, W., Naik, S., Sones, J. E., & Zakharov, E. V. (2018). A Sequel to Sanger: amplicon sequencing that scales. BMC Genomics, 19(1), 219. 10.1186/s12864-018-4611-3

12. Hebert, P. D. N., Floyd, R., Jafarpour, S., & Prosser, S. W. J. (2025). Barcode 100K specimens: In a single nanopore run. Molecular Ecology Resources, 25(1), e14028. 10.1111/1755-0998.14028

13. Hebert, P. D. N., Ratnasingham, S., & DeWaard, J. R. (2003). Barcoding animal life: cytochrome c oxidase subunit 1 divergences among closely related species. Proceedings of the Royal Society B: Biological Sciences, 270(*Suppl. 1*), S96–S99. 10.1098/rsbl.2003.0025

14. Ivanova, N. V., deWaard, J. R., & Hebert, P. D. N. (2006). An inexpensive, automation-friendly protocol for recovering high-quality DNA. Molecular Ecology Notes, 6(4), 998–1002. 10.1111/j.1471-8286.2006.01428.x

15. Jain, M., Olsen, H. E., Paten, B., & Akeson, M. (2016). The Oxford Nanopore MinION: delivery of nanopore sequencing to the genomics community. Genome Biology, 17(1), 239. 10.1186/s13059-016-1103-0

16. Jenjaroenpun, P., Wongsurawat, T., Wadley, T. D., Wassenaar, T. M., Liu, J., Dai, Q., Wanchai, V., Akel, N. S., Jamshidi-Parsian, A., Franco, A. T., Boysen, G., Jennings, M. L., Ussery, D. W., He, C., & Nookaew, I. (2021). Decoding the epitranscriptional landscape from native RNA sequences. Nucleic Acids Research, 49(2), e7. 10.1093/nar/gkaa620

17. Katoh, K., & Standley, D. M. (2013). MAFFT multiple sequence alignment software version 7: improvements in performance and usability. Molecular Biology and Evolution, 30(4), 772–780. 10.1093/molbev/mst010

18. Koblmüller, S., Resl, P., Klar, N., Bauer, H., Zangl, L., & Hahn, C. (2024). DNA barcoding for species identification of moss-dwelling invertebrates: Performance of nanopore sequencing and coverage in reference database. Diversity, 16(4), 196. 10.3390/d16040196

19. Kokoris, M., McRuer, R., Nabavi, M., Jacobs, A., Prindle, M., Cech, C., Berg, K., Lehmann, T., Machacek, C., Tabone, J., Chandrasekar, J., McGee, L., Lopez, M., Reid, T., Williams, C., Barrett, S., Lehmann, A., Kovarik, M., Busam, R., … Corning, M. (2025). Sequencing by Expansion (SBX) - a novel, high-throughput single-molecule sequencing technology. bioRxiv: The Preprint Server for Biology. 10.1101/2025.02.19.639056

20. Li, H. (2018). Minimap2: pairwise alignment for nucleotide sequences. Bioinformatics, 34(18), 3094–3100. 10.1093/bioinformatics/bty191

21. Liu, T., & Conesa, A. (2025). Profiling the epigenome using long-read sequencing. Nature Genetics, 57(1), 27–41. 10.1038/s41588-024-02038-5

22. Martin, M. (2011). Cutadapt removes adapter sequences from high-throughput sequencing reads. EMBnet.Journal, 17(1), 10. 10.14806/ej.17.1.200

23. Meyer, D., Goettsch, W., Spangenberg, J., Stieber, B., Krautwurst, S., Hölzer, M., Brandt, C., Linde, J., Höner Zu Siederdissen, C., Srivastava, A., Zarkovic, M., Wollny, D., & Marz, M. (2026). Unlocking the full potential of nanopore sequencing: tips, tricks, and advanced data analysis techniques. Nucleic Acids Research, 54(3). 10.1093/nar/gkag023

24. Morinière, J., Balke, M., Doczkal, D., Geiger, M. F., Hardulak, L. A., Haszprunar, G., Hausmann, A., Hendrich, L., Regalado, L., Rulik, B., Schmidt, S., Wägele, J.-W., & Hebert, P. D. N. (2019). A DNA barcode library for 5,200 German flies and midges (Insecta: Diptera) and its implications for metabarcoding-based biomonitoring. Molecular Ecology Resources, 19(4), 900–928. 10.1111/1755-0998.13022

25. Pomerantz, A., Peñafiel, N., Arteaga, A., Bustamante, L., Pichardo, F., Coloma, L. A., Barrio-Amorós, C. L., Salazar-Valenzuela, D., & Prost, S. (2018). Real- time DNA barcoding in a rainforest using nanopore sequencing: opportunities for rapid biodiversity assessments and local capacity building. GigaScience, 7(4), giy033. 10.1093/gigascience/giy033

26. Prosser, S. W. J., Floyd, R. M., Thompson, K. A., Monckton, S. K., & Hebert, P. D. N. (2025). BOLDistilled: Automated construction of comprehensive but compact DNA barcode reference libraries. Molecular Ecology Resources, 25(8), e70043. 10.1111/1755-0998.70043

27. Prosser, S. W. J., Thompson, K. A., Bard, N. W., Floyd, R. M., Ozsahin, E., & Hebert, P. D. N. (2026). The Barcode Inference Pipeline (BIP): From Sequencer Output to DNA Barcodes. bioRxiv. 10.64898/2026.07.21.739876

28. Pugh, J. (2023). The current state of nanopore sequencing. Methods in Molecular Biology, 2632, 3–14. 10.1007/978-1-0716-2996-3_1

29. R Core Team. (2024). R: A Language and Environment for Statistical Computing. R Foundation for Statistical Computing, Vienna, Austria. https://www.R-project.org

30. Ratnasingham, S., & Hebert, P. D. N. (2013). A DNA-based registry for all animal species: the Barcode Index Number (BIN) system. PloS One, 8(7), e66213. 10.1371/journal.pone.0066213

31. Ratnasingham, S., Wei, C., Chan, D., Agda, J., Agda, J., Ballesteros-Mejia, L., Boutou, H. A., El Bastami, Z. M., Ma, E., Manjunath, R., Rea, D., Ho, C., Telfer, A., McKeowan, J., Rahulan, M., Steinke, C., Dorsheimer, J., Milton, M., & Hebert, P. D. N. (2024). BOLD v4: A centralized bioinformatics platform for DNA-based biodiversity data. Methods in Molecular Biology (Clifton, N.J.), 2744, 403–441. 10.1007/978-1-0716-3581-0_26

32. Rognes, T., Flouri, T., Nichols, B., Quince, C., & Mahé, F. (2016). VSEARCH: a versatile open source tool for metagenomics. PeerJ, 4, e2584. 10.7717/peerj.2584

33. Schoch, C. L., Seifert, K. A., Huhndorf, S., Robert, V., Spouge, J. L., Levesque, C. A., & Chen, W. (2012). Nuclear ribosomal internal transcribed spacer (ITS) region as a universal DNA barcode marker for Fungi. Proceedings of the National Academy of Sciences of the United States of America, 109(*16*), 6241–6246. 10.1073/pnas.1117018109

34. Silvestre-Ryan, J., & Holmes, I. (2021). Pair consensus decoding improves accuracy of neural network basecallers for nanopore sequencing. Genome Biology, 22(1), 38. 10.1186/s13059-020-02255-1

35. Song, H., Liu, J., Yang, F., Du, W., Qin, L., Su, Y., Jiang, L., & Wang, Z. (2025). A long-amplicon nanopore sequencing and analysis method for human whole mitochondrial genome. Forensic Science International, 377, 112668. 10.1016/j.forsciint.2025.112668

36. Srivathsan, A., Baloğlu, B., Wang, W., Tan, W. X., Bertrand, D., Ng, A. H. Q., Boey, E. J. H., Koh, J. J. Y., Nagarajan, N., & Meier, R. (2018). A MinION^TM^-based pipeline for fast and cost-effective DNA barcoding. Molecular Ecology Resources, 18(5), 1035–1049. 10.1111/1755-0998.12890

37. Srivathsan, A., Feng, V., Suárez, D., Emerson, B., & Meier, R. (2024). ONTbarcoder 2.0: rapid species discovery and identification with real-time barcoding facilitated by Oxford Nanopore R10.4. Cladistics: The International Journal of the Willi Hennig Society, 40(2), 192–203. 10.1111/cla.12566

38. Srivathsan, A., Lee, L., Katoh, K., Hartop, E., Kutty, S. N., Wong, J., Yeo, D., & Meier, R. (2021). ONTbarcoder and MinION barcodes aid biodiversity discovery and identification by everyone, for everyone. BMC Biology, 19(1), 217. 10.1186/s12915-021-01141-x

39. Whiteford, N. (2022a). Thoughts on Chinese Nanopore Startups. https://aseq.substack.com/p/thoughts-on-chinese-nanopore-startups

40. Whiteford, N. (2022b, May 11). Qitan Technology. ASeq Newsletter. https://aseqer.com/p/qitan-technology

41. Wickham, H., Averick, M., Bryan, J., Chang, W., McGowan, L., François, R., Grolemund, G., Hayes, A., Henry, L., Hester, J., Kuhn, M., Pedersen, T., Miller, E., Bache, S., Müller, K., Ooms, J., Robinson, D., Seidel, D., Spinu, V., … Yutani, H. (2019). Welcome to the Tidyverse. Journal of Open Source Software, 4(43), 1686. 10.21105/joss.01686

42. Yu, D. W., Ji, Y., Emerson, B. C., Wang, X., Ye, C., Yang, C., & Ding, Z. (2012). Biodiversity soup: metabarcoding of arthropods for rapid biodiversity assessment and biomonitoring. Methods in Ecology and Evolution, 3(4), 613–623. 10.1111/j.2041-210x.2012.00198.x

43. Zhang, T., Li, H., Ma, S., Cao, J., Liao, H., Huang, Q., & Chen, W. (2023). The newest Oxford Nanopore R10.4.1 full-length 16S rRNA sequencing enables the accurate resolution of species-level microbial community profiling. Applied and Environmental Microbiology, 89(10), e0060523. 10.1128/aem.00605-23

44. Zhao, M., Song, H., Zheng, Y., Qin, L., Liu, X., Deng, S., Liu, J., Chen, L., Du, W., & Wang, Z. (2026). A proof-of-principle study: Species identification based on mitochondrial 12S rRNA gene using QNome nanopore sequencing. Forensic Science International. Genetics, 81, 103388. 10.1016/j.fsigen.2025.103388

